# Activation of ChvG-ChvI regulon by cell wall stress confers resistance to β-lactam antibiotics and initiates surface spreading in *Agrobacterium tumefaciens*

**DOI:** 10.1101/2022.05.26.493563

**Authors:** Michelle A. Williams, Jacob M. Bouchier, Amara K. Mason, Pamela J.B. Brown

**Author notes:** denotes equal contributions (Michelle A. Williams, Jacob M. Bouchier). Boulevard Brewery, 2501 Southwest Blvd, Kansas City, MO 64108.

## Abstract

A core component of nearly all bacteria, the cell wall is an ideal target for broad spectrum antibiotics. Many bacteria have evolved strategies to sense and respond to antibiotics targeting cell wall synthesis, especially in the soil where antibiotic-producing bacteria compete with one another. Here we show that cell wall stress caused by both chemical and genetic inhibition of the essential, bifunctional penicillin-binding protein PBP1a prevents microcolony formation and activates the canonical host-invasion two-component system ChvG-ChvI in *Agrobacterium tumefaciens*. Using RNA-seq, we show that depletion of PBP1a for 6 hours results in a downregulation in transcription of flagellum-dependent motility genes and an upregulation in transcription of type VI secretion and succinoglycan biosynthesis genes, a hallmark of the ChvG-ChvI regulon. Depletion of PBP1a for 16 hours, results in differential expression of many additional genes and may promote a general stress response. Remarkably, the overproduction of succinoglycan causes cell spreading and deletion of the succinoglycan biosynthesis gene *exoA* restores microcolony formation. Treatment with cefsulodin phenocopies depletion of PBP1a and we correspondingly find that *chvG* and *chvI* mutants are hypersensitive to cefsulodin. This hypersensitivity only occurs in response to treatment with β-lactam antibiotics and moenomycin, suggesting that the ChvG-ChvI pathway may play a key role in resistance to antibiotics targeting cell wall synthesis. Finally, we provide evidence that ChvG-ChvI likely has a conserved role in conferring resistance to cell wall stress within the Alphaproteobacteria.

**AUTHOR SUMMARY:** Soil dwelling bacteria reside in changing environments requiring them to frequently adapt to stressful conditions to ensure survival. The bacterial envelope provides structural integrity and protection against osmotic stress and turgor pressure imposed by the environment. While the mechanisms of cell membrane and cell wall biogenesis have been extensively studied, our understanding of how diverse microbes respond to cell envelope and cell wall stress to increase their fitness remains limited. In this work, we identify ChvG-ChvI regulon as an envelope stress response system that confers protection under cell wall stress conditions in the bacterial plant pathogen *Agrobacterium tumefaciens*. This is a new function for the well-characterized ChvG-ChvI pathway which is also acid induced and promotes plant host invasion. Our results suggest that the ChvG-ChvI pathway has a broadly conserved role in protecting Alphaproteobacterial cells from extracellular stress and a more specific role in response to acid stress and promoting plant-microbe interactions.

## INTRODUCTION

The soil environment is constantly in flux and can undergo rapid changes in hydration, nutrient availability, temperature, acidity levels and many other abiotic and biotic factors [1]. To survive in these conditions, soil-dwelling bacteria must be able to monitor and respond to the changes around them. One of the main mechanisms bacteria employ to monitor changes in their environment is coupling environmental stimuli to transcriptional regulation using two-component systems (TCS) [2]. In turn transcriptional changes can modify bacterial behavior. In the plant-pathogen *Agrobacterium tumefaciens*, two TCS sense the presence of a potential host and initiate transcription programs that transition the bacterium into a virulent state [3].

The VirA-VirG histidine kinase/response-regulator pair is a characteristic of Agrobacteria and responds to plant phenolic compounds such as acetosyringone. Activation induces expression of the *vir* regulon, which encodes genes that are required for pathogenicity and plant transformation [4]. The ChvG-ChvI TCS is more broadly conserved across many Alphaproteobacteria but has been best characterized among the plant symbionts of Rhizobiales such as *Sinorhizobium meliloti* [5,6]. Activation of ChvG-ChvI is required for the transition from a free-living bacterium to a host-associated lifestyle [7].

In *S. meliloti* and *A. tumefaciens* ChvG-ChvI is regulated by the periplasmic protein ExoR. Under neutral conditions, ExoR binds to and represses ChvG; however, when cells are exposed to acidic conditions, ExoR is proteolyzed, which allows for activation of the ChvG-ChvI TCS [8,9]. ChvI induces transcriptional changes in many genes across several major pathways. For example, ChvI upregulates transcription of *mirA*, encoding a repressor of the motility response regulator Rem and ultimately resulting in suppression of genes for motility and chemotaxis [10]. ChvI also upregulates genes for exopolysaccharide production and, in *A. tumefaciens*, induction of the Type VI Secretion System (T6SS) [11].

Conservation of the ChvG-ChvI TCS is taxonomically constrained to several orders of Alphaproteobacteria, many of whom have free-living lifestyles that are never host-associated [6]. This begs the question: why is the ChvG-ChvI pathway conserved in so many non-host-associated bacteria? Recent interest in the ChvG-ChvI pathway of *Caulobacter crescentus* (ChvGI) provides a glimpse at the function of the pathway in the context of a bacterium with a drastically different ecological niche to that of *A. tumefaciens* or *S. meliloti*. ChvGI of *C. crescentus* senses and responds to osmotic stress and mutants of ChvGI are sensitive to several cell-wall targeting antibiotics [12,13]. It remains unclear if this function is solely a characteristic of *C. crescentus* ChvGI or if it is conserved across ChvG-ChvI orthologs.

Although the cell wall is an essential feature of bacteria that protects them from environmental stressors, relatively little is known about how bacteria sense and respond to changes in the composition of their cell wall. Peptidoglycan (PG) is a heteroglycan decorated with cross-linked peptide stems and is the primary component of bacterial cell walls. During elongation in *A. tumefaciens*, nascent PG insertion is constrained to the pole. Polar growth is a characteristic of Rhizobiales and does not require the canonical MreB-RodA-PBP2 elongation complex. Indeed, all members of Rhizobiales have lost this complex entirely [14,15].

We showed that PBP1a is essential in *A. tumefaciens* and is the primary driver of polar growth. Depletion of PBP1a eliminates nascent PG insertion at the growth pole, leading to shorter cells that have compositional changes in PG [16]. In addition to its role in polar PG insertion, here we observe that during PBP1a depletion cells spread apart rather than form microcolonies. To better understand this phenotype, we used RNA-seq to obtain transcriptional profiles of cells depleted of PBP1a after 6 hours, corresponding to the onset of the spreading phenotype, and after 16 hours. Transcriptomic changes closely mimic the transcriptome changes seen when ChvG-ChvI is activated in *A. tumefaciens*, including downregulation of genes for motility and chemotaxis and upregulation of genes for exopolysaccharide biosynthesis and T6SS. Here we experimentally validate the RNA-seq results, confirming the impacts of PBP1a depletion on the physiology and behavior of *A. tumefaciens*.

## RESULTS

### PBP1a depletion prevents proper microcolony formation

Here, we grew PBP1a depleted cells on agarose pads and saw that these cells exhibit surface spreading rather than forming closely packed microcolonies (Fig 1A). Additionally, when centrifuging cultures of PBP1a-depleted cells, we observed that the cells did not pellet (Fig S1). Considering the possibility that depletion of PBP1a somehow signals for these phenomena, we decided to look at RgsM, another enzyme required for polar elongation. Previous work points to RgsM activity being required for incorporation of nascent PG by PBP1a [17]. However, depletion of RgsM did not cause surface spreading (Fig 1A) indicating that an imbalance of PG hydrolysis and synthesis triggers spreading and the inability pellet in *A. tumefaciens*. Deletions of genes encoding other high molecular weight PBPs and *mtgA*, a PG transglycosylase did not induce spreading (Fig 1B).

**Fig 1.**
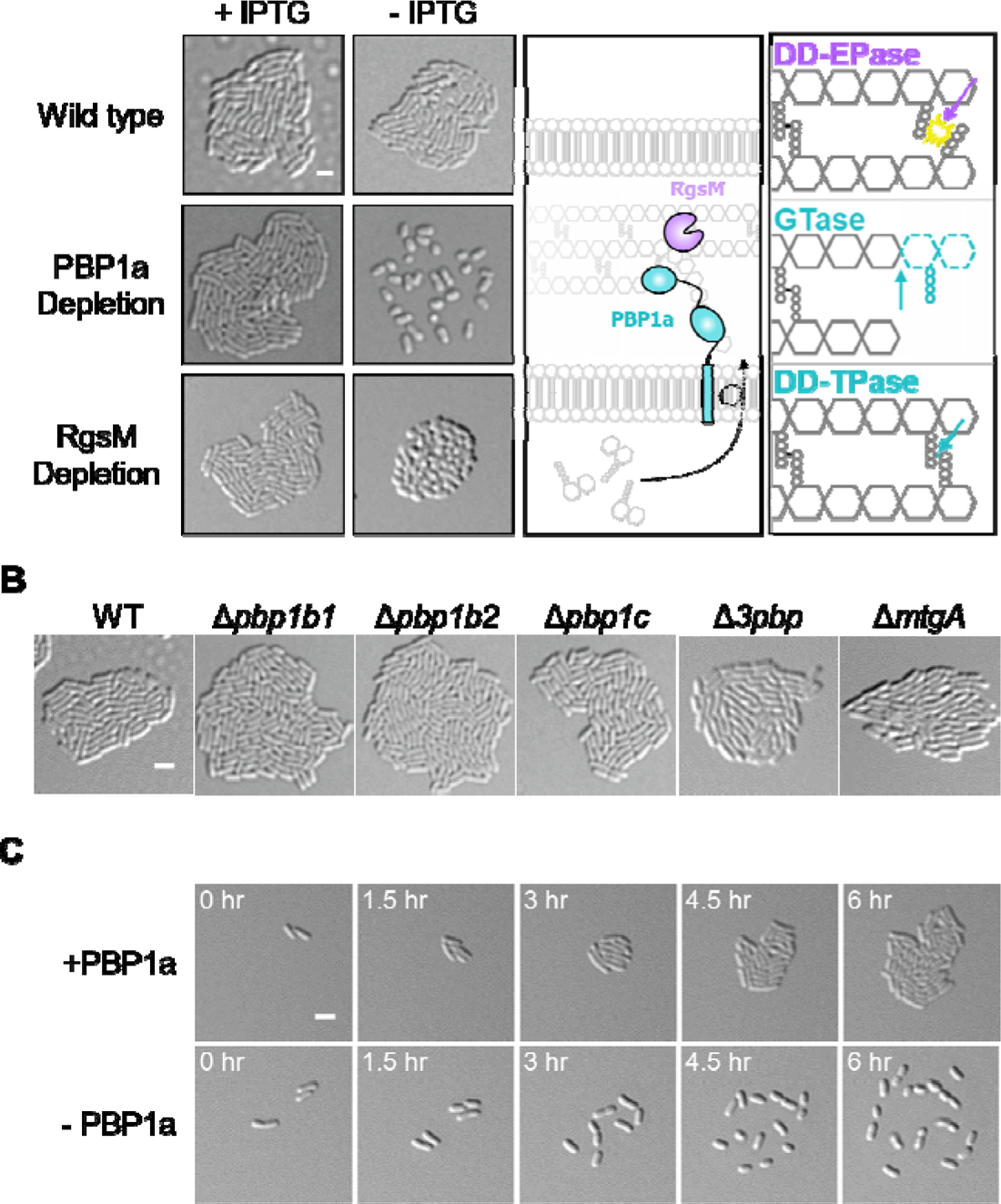
The PBP1a depletion fails to form microcolonies independent of flagellar motility. A. Micrographs of wildtype, PBP1a depletion, and RgsM depletion with or without 1mM IPTG inducer. Each strain was grown to exponential phase, spotted on an ATGN agar pad, allowed to grow for 16 hours, and imaged by DIC microscopy. Scale bar depicts 2μm. The graphic depicts the working model that RgsM _DD_-endopeptidase activity is required for incorporation of nascent glycan strands into the preexisting peptidoglycan (PG) macromolecule by PBP1a. RgsM cleaves _DD_-crosslinks, PBP1a glycosyltransferase activity incorporates lipid II into the PG glycan strand, PBP1a _DD_-transpeptidase activity crosslinks the peptide stem of the nascent PG, fully incorporating it into the macromolecule. EPase, endopeptidase; GTase, glycosyltransferase; TPase, transpeptidase. B. Micrographs of wild type, Δ*pbp1b1*, Δ*pbp1b2*, Δ*pbp1c*, and Δ*mtgA*. Each strain was grown to exponential phase, spotted on an ATGN agar pad, allowed to grow for 16 hours, and imaged by DIC microscopy. Scale bar depicts 2μm. C. Time-lapse microscopy of the PBP1a depletion grown on an agar pad with or without 1mM IPTG inducer. DIC images were acquired every 10 minutes. Time is shown in hours. For the - PBP1a strain, cells were washed 3X with ATGN media and grown at 28 C with shaking for 4 hours before cells were spotted on an agar pad for imaging.

Timelapse microscopy revealed that after ∼6 hours of PBP1a depletion cells spread apart, though the movement of cell appears to be confined within a relatively small region of the agarose pad (Fig 1C, Movie S1). Since spreading is confined and occurs over the course of many hours, we suspected that this phenomenon was not simply caused by the activation of swimming motility.

### PBP1a depletion induces global transcriptome changes

To understand the spreading phenotype caused by PBP1a depletion, we compared the transcriptomes of cells at the onset or late stage of the surface spreading phenotype. Cells were grown with or without the inducer Isopropyl β-D-1-thiogalactopyranoside (IPTG) for *mrcA*, encoding PBP1a, expression for 6 or 16 hours (Fig. 2A). As a baseline, we compared transcriptional profiles of WT in the presence and absence of IPTG to the PBP1a depletion strain in the presence of IPTG. The addition of IPTG did not alter gene expression profiles of WT cells, and only minor differences were apparent between the PBP1a depletion strain background and WT when both strains are grown in the presence of IPTG (Fig S2). We next compared differences in the PBP1a replete strain to the PBP1a depleted strain at either 6- or 16-hours post depletion (Fig 2A). Using a p-value < 0.05 and log2 fold-change (L2FC) > 2.0, we identified 91 and 306 genes that were differentially expressed in the + PBP1a strain compared to the 6- or 16-hour depletion, corresponding to 2% and 6% of the total genes, respectively.

**Fig 2.**
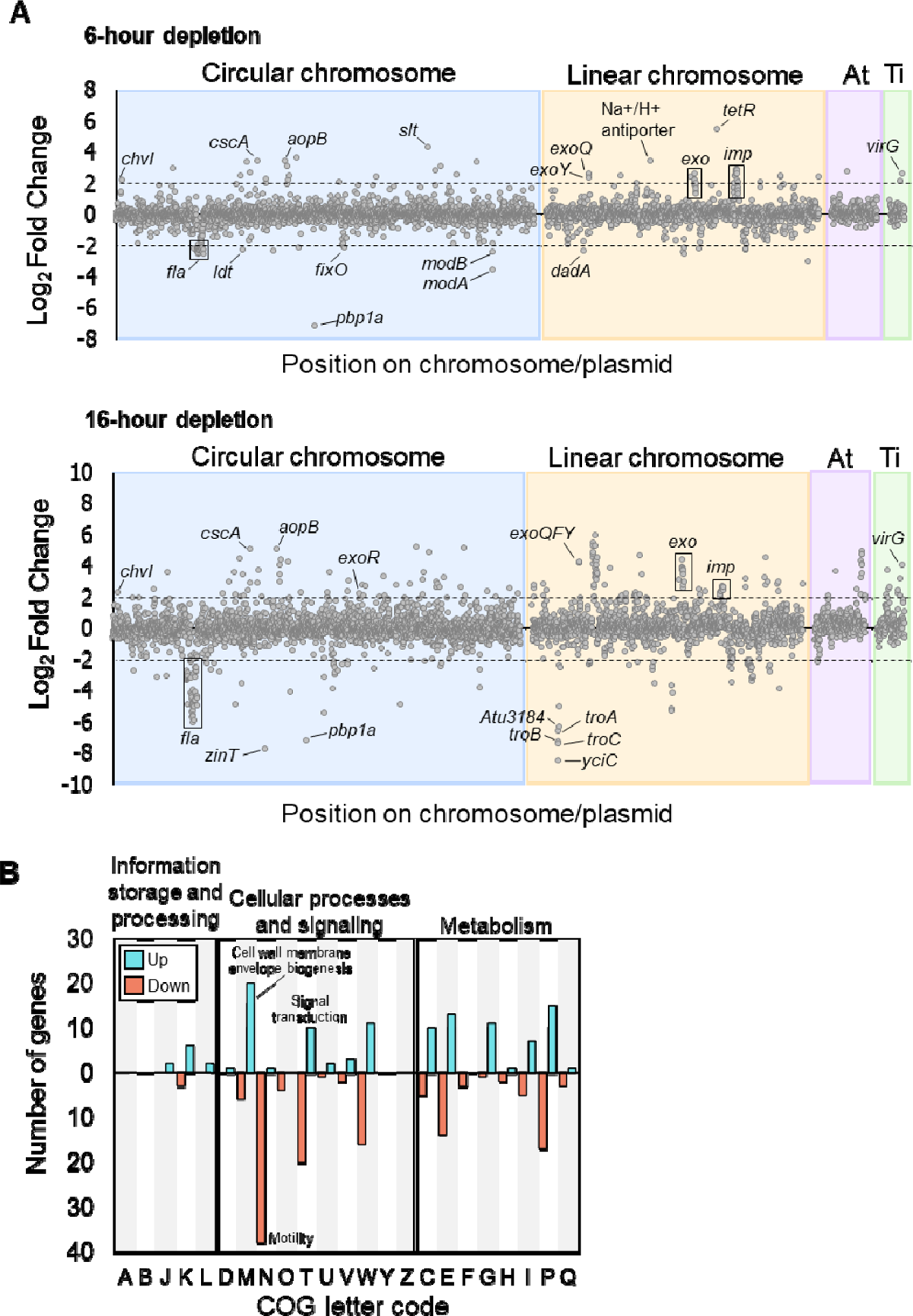
Analysis of the PBP1a depletion transcriptomes by RNA-seq. A. Plots comparing Log2Fold Change of the + PBP1a transcriptome to that of the - PBP1a 6-hour transcriptome and to that of the 16-hour depletion. Gray dots represent a single transcript, and the dotted lines represent +/-2.0 Log2Fold Change threshold. Plots are delimited by chromosome and mega plasmid. B. COG categorical analysis of the 16-hour depletion of PBP1a. Pink, downregulated; Cyan, upregulated.

Overall, we observed large-scale changes in a diverse and widespread range of genes that are regulated in response to PBP1a depletion. Initially, the response to PBP1a depletion is primarily mediated by chromosomally encoded rather than plasmid encoded genes. *A. tumefaciens* has a circular chromosome, which houses roughly half (51.7%) of the protein-coding genes, a linear chromosome (34.7%) and two mega plasmids, the At plasmid (10%) and Ti plasmid (3.6%). Most of the genes differentially expressed at both time points during PBP1a depletion were encoded on the linear and circular chromosomes (Fig. 2A). Most of the differentially abundant genes from the 6-hour timepoint were also present in the 16-hour timepoint. For several of these genes, the magnitude of differential transcript abundance remained relatively constant. For example, the response regulator ChvI, had an increased relative abundance early in response to PBP1a depletion that remained constant in the 16-hour timepoint. In contrast, several genes displayed a continuous increase or decrease in transcript abundance between the 6- and 16-hour timepoints, including genes that encode proteins necessary for assembly of flagella and type 6 secretion system machinery. Finally, several genes were only differentially abundant at the 16-hour timepoint, including many genes encoding proteins important for cell envelope homeostasis such as the Tol-Pal system [18], and >30 ABC transporters.

To further categorize the diverse set of differentially abundant genes we identified Clusters of Orthologous Groups (COGs) in the 16-hour timepoint and classified them based on functional categories represented by a single letter code (Figure 2B) [19,20]. The most affected COG category was motility (N). Decreased abundance of mRNAs containing genes which encode structural flagella proteins further supports the hypothesis that the spreading phenotype is independent of flagella-based motility. The COG category with the largest proportion of increased differentially abundant genes was cell wall, membrane, and envelope biogenesis (M). Notably, no significant changes in the transcripts of other penicillin-binding proteins or glycosyltransferases were observed in response to loss of PBP1a (Table S1). However, significant changes in transcripts encoding cell wall remodeling enzymes such as LD-transpeptidases, endopeptidases, and soluble lytic transglycosylases were detected (Table S1). Atu0844, an LD-transpeptidase, was strongly downregulated suggesting it may play an important role in polar growth alongside PBP1a. Additionally, one putative β-lactamase gene, Atu0933, was strongly upregulated, which may provide a strategy to protect against cell wall damage. In addition, many of the genes found in this COG category encode cell envelope homeostasis and osmotic stress response proteins, including the Tol-Pal system, several outer membrane proteins (i.e. AopB), and periplasmic sensors (i.e. CreD).

At the 16-hour timepoint, the largest changes to cellular metabolism occurred in the inorganic ion transport and metabolism (P) and amino acid metabolism and transport (E) COG categories, suggesting a shift in nutrient uptake and metabolism, as expected during a general stress response. Interestingly, the six most downregulated genes in the 16-hour timepoint, with the exception of *mrcA*, encoding PBP1a, were *yciC* (Atu3181), *zinT* (Atu1049), *troC* (Atu3180), *troB* (Atu3179), *troA* (Atu3179), and Atu3184, all of which are major components of cytoplasmic zinc uptake in *Agrobacterium tumefaciens* (Fig 2B, bottom) [21].

We also observed large increases and decreases in the transcript abundance of signal transduction genes. Transcription of *exoR* (Atu1715), *chvG* (Atu0033), and *chvI* (Atu0034) were upregulated at both the 6-hour (L2FC = 0.995, 1.42, and 2.18) and 16-hour (L2FC = 2.20, 1.41, and 2.27) depletions of PBP1a. Transcription of genes encoding three other two-component systems were also strongly upregulated (Fig S3, Table S1).

### Transcriptome changes during PBP1a depletion mimic activation of the ChvG-ChvI two-component system

Transcription of *virG*, encoding a TCS response regulator, was also strongly upregulated in both the 6-hour (L2FC = 2.61) and 16-hour (L2FC = 4.02) timepoints (Fig S3, Table S1). Transcription of *virG* has been reported to be upregulated under both host-invasion and virulence-inducing conditions [11]. Because *virG* was also upregulated during depletion of PBP1a, we reasoned that PBP1a depletion may be mimicking one of these two conditions. Using comparative transcriptomics, we compared the 150 most differentially expressed genes (DEGs) against published datasets that simulate host-invasion conditions (Δ*exoR &* pH 5.5) and virulence-inducing conditions (acetosyringone treatment & growth on AB media) [22][23]. We found that L2FC values of the 6-hour PBP1a depletion RNA-seq dataset correlated with the two host-invasion conditions and not with the virulence-inducing datasets, as indicated by the spearmen rho correlation coefficient for each comparison (Δ*exoR*, rho = 0.875; pH 5.5, rho = 0.766) (Fig 3 and S4). Rho values near 1 indicate similar DEGs between each dataset. Rho values near 0 would indicate no similar DEGs between each dataset (Fig 3A). Interestingly, each of these two datasets have been implicated in activation of the ChvG-ChvI pathway [9,11]. Correlation with each strongly implicates ChvG-ChvI activation in our RNA-seq, suggesting that depletion of PBP1a may provide a signal leading to similar changes described to occur during the transition to a host-invasion lifestyle. While this trend was maintained in the 16-hour timepoint, we observed additional genes that were differentially expressed under depletion of PBP1a, but not in the Δ*exoR* and pH 5.5 datasets (Fig 3, Fig S4B). Indeed, the rho values for the 16-hour depletion of PBP1a compared to the host-invasion datasets (Δ*exoR*, rho = 0.529; pH 5.5, rho = 0.739 were lower than the 6-hour comparisons. Additionally, we found 217 more genes with L2FC > 2.0 in the 16-hour depletion than in the 6-hour depletion (Fig 3B). Together, these findings suggest that longer depletions of PBP1a may result in the activation of additional regulons beyond ChvG-ChvI.

**Fig 3.**
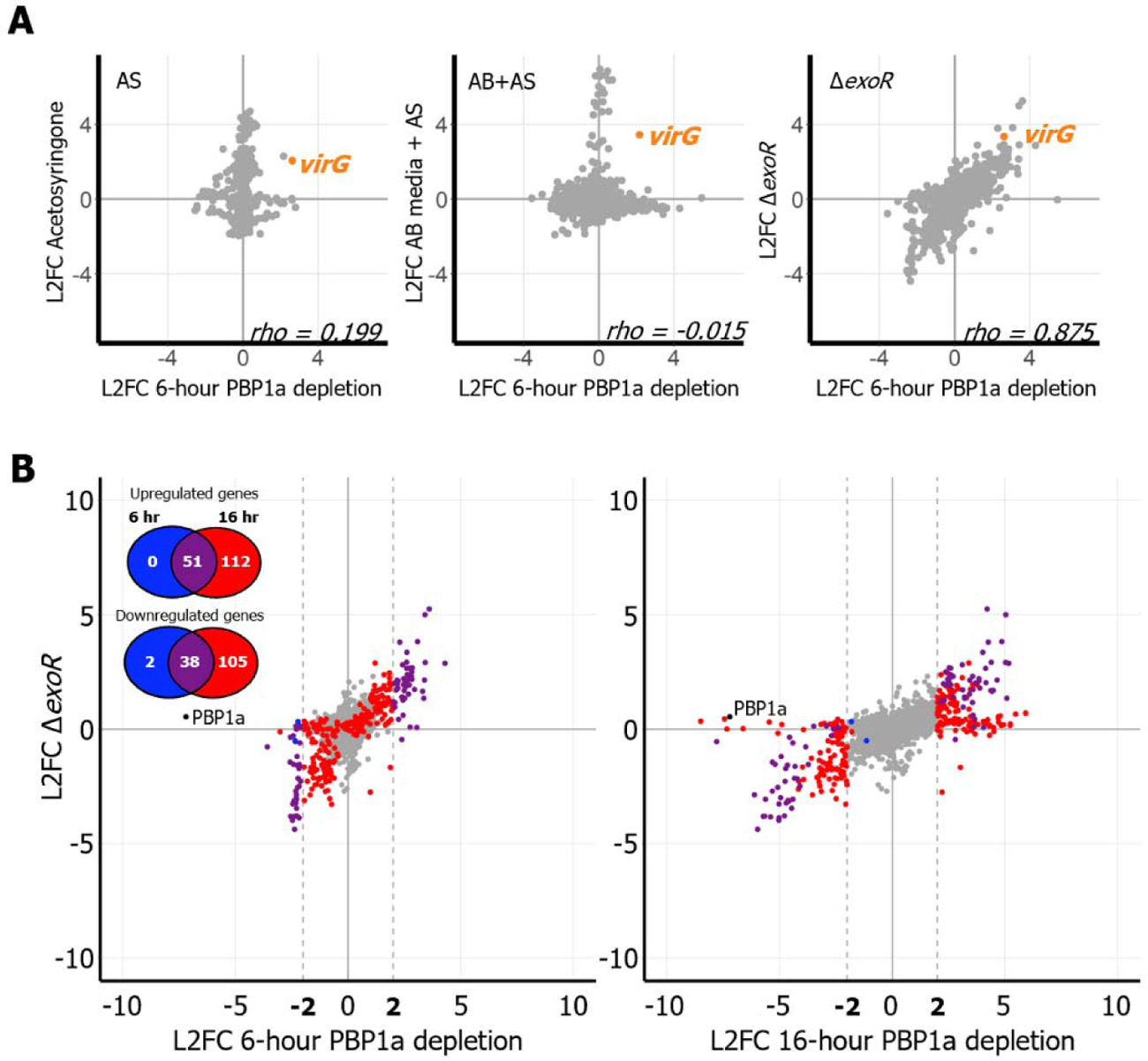
The response to the depletion of PBP1a mimics transcriptional changes associated with host invasion. A. Correlation scatterplots depicting relationships between the log2fold-change (L2FC) values in the 6-hour PBP1a depletion and transcriptomic data sets taken under simulated virulence-inducing conditions (AS and AB+AS) and under simulated host-invading conditions (Δ*exoR*). Each point represents a unique transcript. AS, acetosyrinogone; AB, *Agrobacterium* minimal media; rho, Spearman correlation coefficient. B. Correlation scatterplots comparing L2FC values of transcripts in the Δ*exoR* microarray to either the 6-hour or 16-hour PBP1a depletion. Each transcript is colored according to its change in L2FC values from 6 hours of PBP1a depletion to 16 hours of depletion. Gray, no change; Blue, transcript has |L2FC| > 2.0 in the 6-hour but not in the 16-hour depletion; Red, |L2FC| > 2.0 in the 16-hour but not in the 6-hour depletion; Purple, |L2FC| > 2.0 in both the 6-hour and 16-hour depletion.

Overall, a large number and variety of genes are regulated in response to depletion of PBP1a. Although many of these changes in gene expression have been reported previously in response to low pH or deletion of the ChvG-ChvI negative regulator ExoR, these changes have never been associated with loss of a cell wall synthase in *A. tumefaciens*. These observations indicate that there are additional mechanisms that can activate the ChvG-ChvI TCS.

### Succinoglycan overproduction is required for cell spreading

Previous work has clearly associated activation of ChvG-ChvI to a specific transcriptomic pattern involving downregulation of flagellar motility genes and upregulation of T6SS and succinoglycan biosynthesis genes [9,11,24]. Indeed, this same pattern was observed in the 6- and 16-hour PBP1a depletion datasets (Fig 4A). To confirm that the spreading phenotype is unrelated to flagella-dependent motility, we made an in-frame deletion of *rem*, which encodes a transcriptional regulator of genes encoding structural flagella proteins [25,26], in the PBP1a depletion strain. Deletion of *rem* prevents swimming in *A. tumefaciens* and does not impact microcolony formation on agarose pads (Fig 4B). Upon depletion of PBP1a, *rem* mutants continued to spread, suggesting that the cause of this phenotype is independent of flagella-mediated swimming motility (Fig 4B).

**Fig 4.**
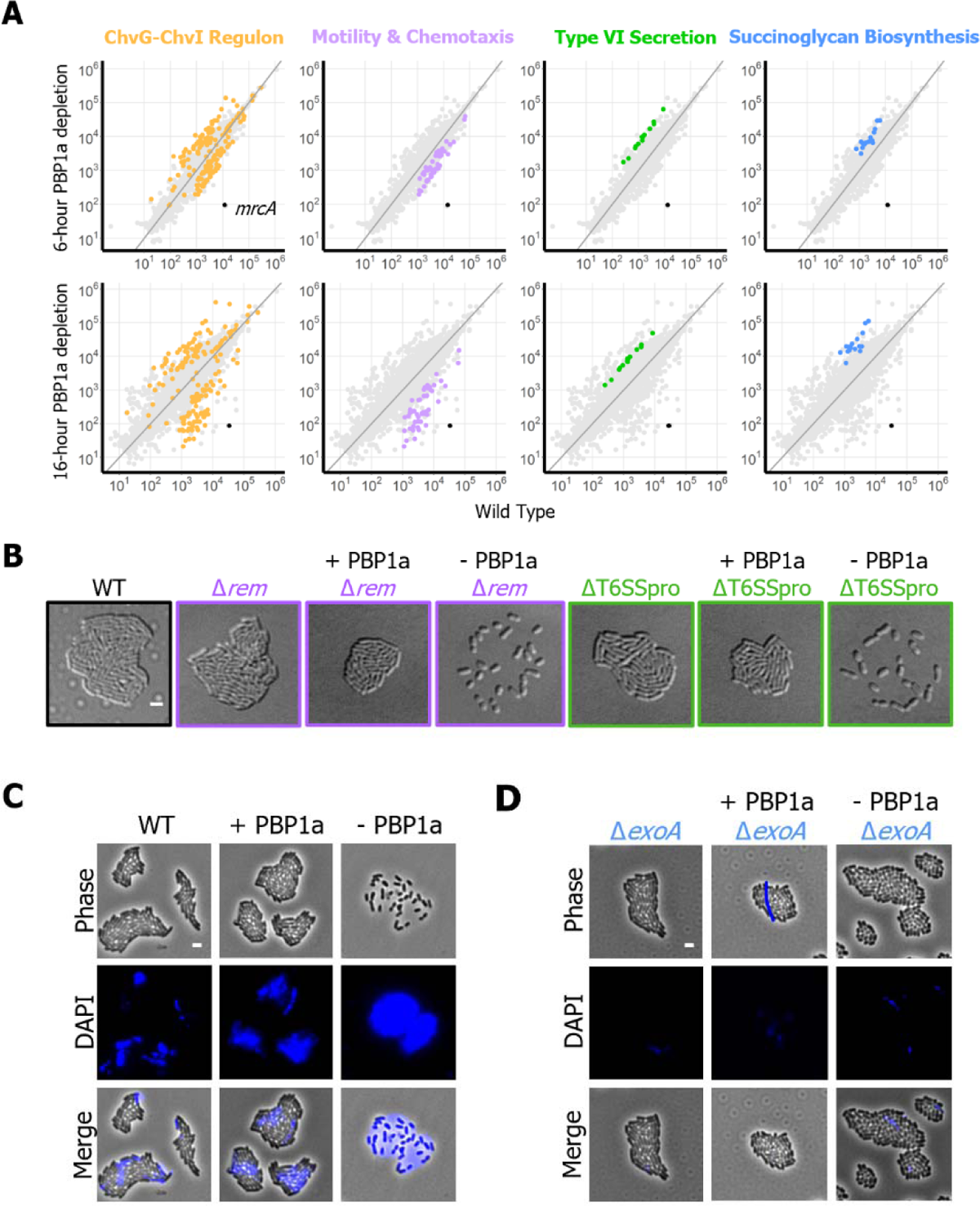
Succinoglycan overproduction is a conserved response to PBP1a depletion and results in failed microcolony formation. A. Scatter plots depicting RPKM values of the 6-hour and 16-hour compared to wild type. Each point represents a unique transcript. Points are colored by category. Gold, ChvG-ChvI regulon; Lavender, Motility and Chemotaxis; Green, Type VI Secretion; Blue, Succinoglycan Biosynthesis; Black, *mrcA* (encoding PBP1a). B. Micrographs of wild type, Δ*rem*, PBP1a replete Δ*rem*, PBP1a depleted Δ*rem*, ΔT6SSpro, PBP1a replete ΔT6SSpro, and PBP1a depleted ΔT6SSpro. Each strain was grown to exponential phase, spotted on a 1% ATGN agar pad containing 1mM IPTG if inducing *mrcA*, allowed to grow for 16 hours, and imaged by DIC microscopy. The scale bar depicts 2μm. C. Micrographs of wild type, and PBP1a depletion with or without IPTG inducer. Each strain was grown to exponential phase and spotted on a 1% ATGN agar pad containing 25μg/mL calcofluor white and 1mM IPTG if inducing *mrcA*. Each was allowed to grow for 16 hours and imaged by phase microscopy with and without the DAPI filter for visualizing calcofluor-stained succinoglycan. D. Micrographs of Δ*exoA* and PBP1a depletion Δ*exoA*, with or without IPTG inducer. Strains were grown and imaged as described for panel C.

All genes in the *imp* and *hcp* operons, which are located on the linear chromosome and encode the structural and toxin proteins of T6SS respectively, are upregulated at both timepoints (Fig 4A). In *A. tumefaciens*, activation of the T6SS results in the production of a contractile nanomachine which delivers effector proteins to antagonize and compete with other bacteria [27]. Among agrobacteria, T6SS is activated by different signals, is important during different stages of the lifecycle, and may be used to acquire nutrients [28]. To determine if the T6SS contributes to the spreading response observed during PBP1a depletion we deleted the intergenic gap between the *hcp* and *imp* operons (ΔT6SSpro). This deletion prevented expression of proteins from both the *hcp* and *imp* operons (Fig S5) [29]. Cell spreading persisted in ΔT6SSpro during depletion of PBP1a suggesting that the activation of T6SS is not responsible for this behavior.

Another possibility is that spreading might be caused by sliding motility, where secretion of a surfactant gives the cells a slippery surface to “slide” across. Notably, *S. meliloti* has been reported to undergo entropy-driven surface spreading during the overproduction of succinoglycan [30]. Succinoglycan is a β-1,4-linked sugar made of glucose and galactose, and is the most abundant exopolysaccharide produced by *A. tumefaciens* and related bacteria [31]. Genes associated with the biosynthesis and secretion of succinoglycan were strongly upregulated in both timepoints. To test if entropy-driven surface spreading is causing PBP1a-depleted *A. tumefaciens* cells to spread, we used a microscopy-based assay to observe succinoglycan production in *A. tumefaciens*. Cells were spotted on agarose pads containing calcofluor white and grown overnight, then imaged using the DAPI filter to detect succinoglycan production (Fig 4C). Wild-type *A. tumefaciens* and the PBP1a replete strains secrete some succinoglycan that enriched near the center of the microcolony (Fig 4C). In comparison, depletion of PBP1a triggers secretion of a large quantity of succinoglycan that defines the boundary of where the cells spread. An in-frame deletion of *exoA*, which encodes a glycosyltransferase required for succinoglycan production in *A. tumefaciens* [32], prevents succinoglycan production (Fig 4D). During PBP1a depletion, microcolony formation is restored in the Δ*exoA* mutant (Fig 4D). Together, these data illustrate that succinoglycan is indeed overproduced and contributes to the surface spreading of the PBP1a depletion.

### Deletion of *chvG* or *chvI* results in hypersensitivity to β-lactam antibiotics

Since activation of succinoglycan production is known to be part of the ChvG-ChvI regulon, we next wanted to test if the PBP1a depletion is activating SGN production through the ChvG-ChvI signaling pathway. We made an in-frame deletion of *chvI* in the PBP1a depletion background and found *chvI* mutants repleted with PBP1a appear morphologically wild-type when grown in minimal media (Fig S6A). However, PBP1a-depleted Δ*chvI* cells were extremely sick. Previously we reported that the PBP1a depletion produces viable daughter cells for up to 5-6 generations [16], however, the PBP1a-depleted Δ*chvI* strain was incapable of a single division event. Instead, the cells exhibited growth arrest and eventually cell lysis shortly after depletion initiation (Fig S6A, Movie S2).

To further assess the enhanced sensitivity of Δ*chvI* to PBP1a depletion, we identified an antibiotic that specifically targets PBP1a enzymatic activity. Treatment with cefsulodin at a concentration of 20 μg/mL resulted in a spreading phenotype similar to the PBP1a depletion (Fig 5A). Additionally, treatment with cefsulodin results in short, round cells, further phenocopying the PBP1a depletion (Fig 5B). Together, these data suggest that cefsulodin targets the transpeptidase activity of PBP1a in *Agrobacterium tumefaciens*, similar to reported cefsulodin specificity in *E. coli* [33]. Next, we observed relative cefsulodin sensitivities in WT, Δ*chvG*, Δ*chvI*, and Δ*exoR* strains. Remarkably, we found that while growth of wild-type and Δ*exoR* cells appeared relatively unaffected by treatment with 10μg/mL of cefsulodin, Δ*chvG* and Δ*chvI* cells were hypersensitive (Fig 4C). Next, we targeted the glycosyltransferase activity of PBP1a through treatment with moenomycin [34], and found that moenomycin-treated cells also spread on agarose pads (Fig S6B). These findings suggest that the ChvG-ChvI TCS is essential for growth when the activity of the major PG synthase is inhibited either chemically or genetically.

**Fig 5.**
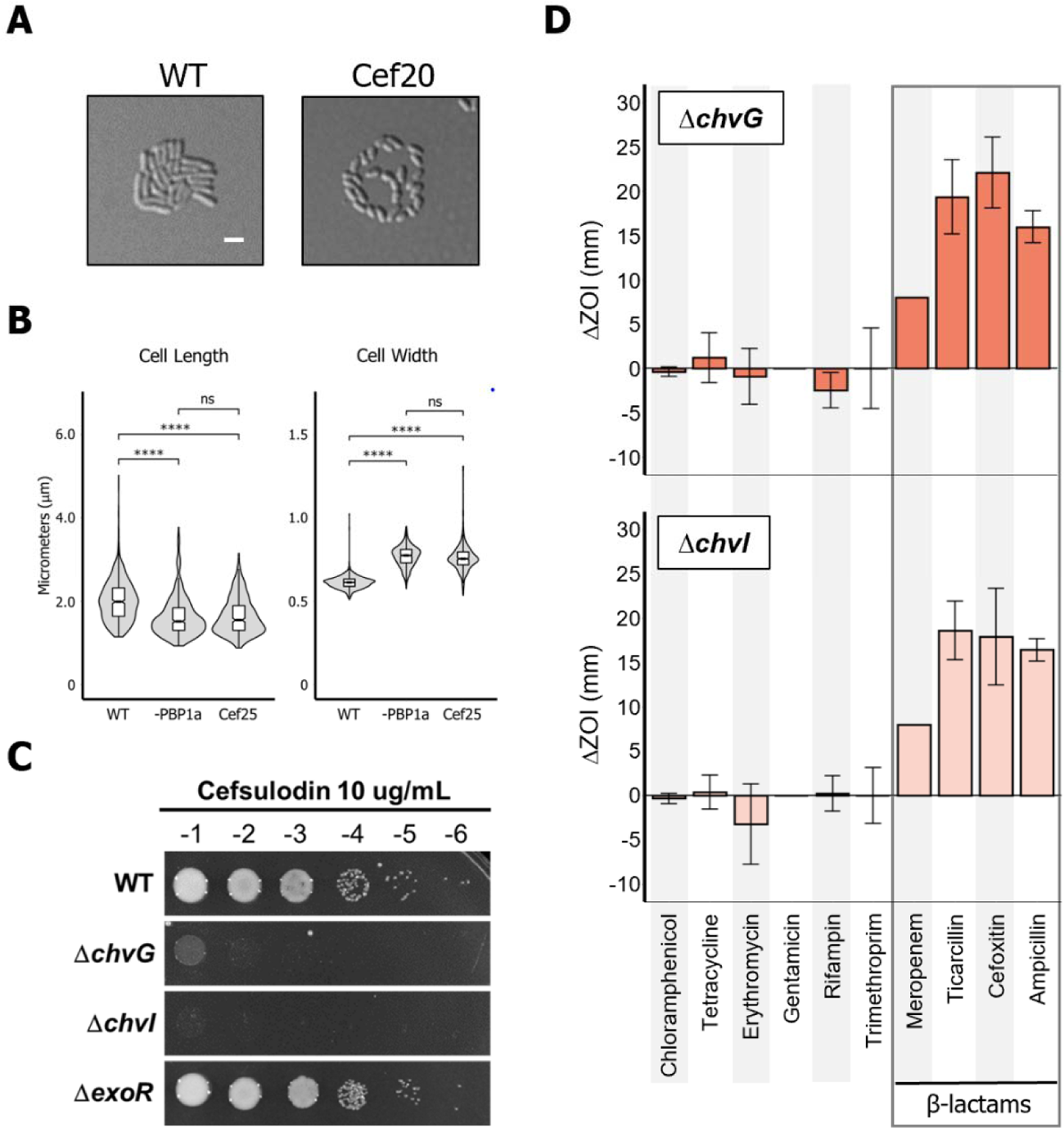
The ChvG-ChvI TCS is conditionally essential under treatment with β-lactam antibiotics. A. Micrographs of untreated and cefsulodin-treated cells. Wild-type cells were grown to exponential phase, spotted on a 1% ATGN agar pad with or without 20 μg/mL cefsulodin and allowed to grow for 16 hours. Each strain was imaged by DIC microscopy. B. Box plots comparing cell length and width between wild-type, PBP1a-depleted, and cefsulodin-treated cells. ns, not significant; ****, p < 0.00005. C. Cell viability of each wild type, Δ*chvG*, Δ*chvI*, and Δ*exoR* spotted on an ATGN agar plate containing 10 μg/mL of cefsulodin. Ten-fold serial dilutions are indicated. D. Graph depicting the change in zone of inhibition from wildtype in Δ*chvG* and Δ*chvI* against ten different antibiotic disks. Error bars represent +/-1 standard deviation from the mean.

We broadened our investigation by testing Δ*chvG* and Δ*chvI* against ten additional antibiotics. four that block protein synthesis (chloramphenicol, tetracycline, erythromycin, and gentamicin); one that blocks DNA replication (nalidixic acid); one that blocks transcription (rifampin); and four other cell wall synthesis inhibiting β-lactam antibiotics (meropenem, cefoxitin, ampicillin, and ticarcillin). To measure changes in sensitivity to each antibiotic compared to wildtype, Δ*chvG* and Δ*chvI* were spread on ATGN minimal media and disks containing each antibiotic were applied. Diameters of the zones of inhibition (ZOI) were measured and the difference in ZOIs for each mutant strain compared to wildtype are shown (Fig 5D). Of the antibiotics tested, Δ*chvG* and Δ*chvI* showed increased sensitivity only to β-lactam antibiotics, suggesting specificity of the ChvG-ChvI pathway in conferring resistance to this antibiotic class.

### ChvG and ChvI are conserved in Alphaproteobacteria but the presence of ExoR is more constrained

The absence of PBP1a activity at the growth pole during elongation activates ChvG-ChvI, the canonical host-invasion pathway of *Agrobacterium tumefaciens*. The ChvG-ChvI pathway is most well known to be activated by environmental changes associated with conditions favorable for plant association, yet this TCS is retained in many non-plant-associated Alphaproteobacteria (Fig 6A). Remarkably, while ChvG-ChvI is conserved in a large proportion of Alphaproteobacteria, ExoR is not (Fig 6A). Predicted structures of the sensor domains of ChvG in bacteria with ChvG-ChvI orthologs show two structural loops (L1 and L2; Fig 6A, Fig S7). While L2 is conserved across the orthologous structures, L1 is expanded solely in the Rhizobiales (Fig S8). This expansion coincides with the retention of ExoR, making it a compelling target for ExoR-ChvG association studies (Fig 6A). The more broadly conserved L2, on the other hand, may enable a conserved, ExoR-independent mechanism for activation of ChvG-ChvI.

**Fig 6.**
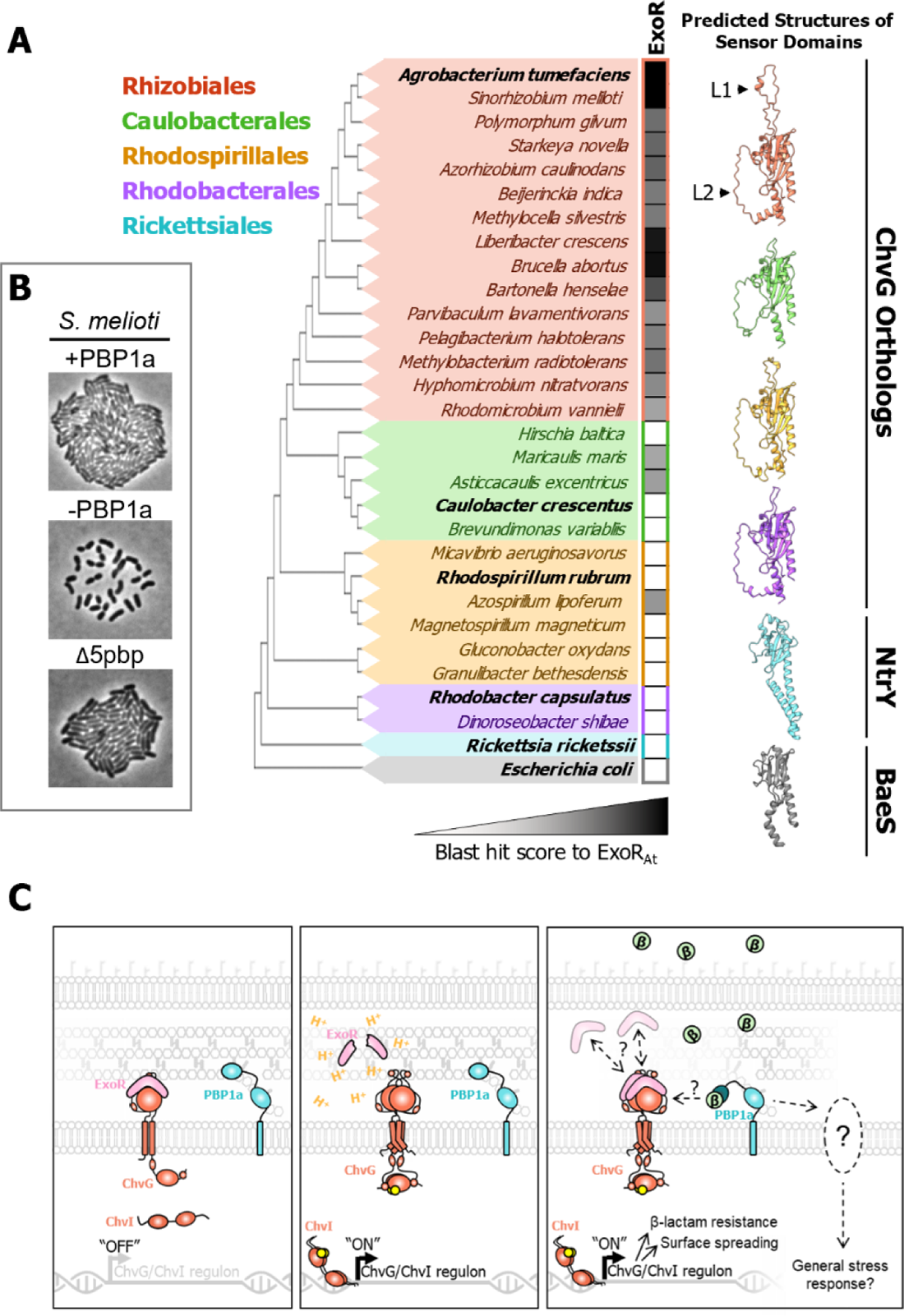
Conservation constraints of ExoR suggest conserved ChvG-ChvI response is independent of ExoR. A. Maximum parsimony tree constructed using MUSCLE sequence alignment [60] on the periplasmic regions of ChvG orthologs. In clades that don’t have a ChvG ortholog, the protein with the highest sequence similarity to ChvG was used instead. Conservation of ExoR was calculated using blast max scores from top hits when protein blasting [59] ExoR from *Agrobacterium tumefaciens* against each species in the tree. Phyre2 predicted structures of periplasmic domains of ChvG orthologs from representatives (bold) in each genus are shown. Conserved structural loops are denoted as L1 and L2. B. Micrographs of *Sinorhizobium meliloti* Rm2011 PBP1a replete, PBP1a depleted, and a strain with deletions of genes encoding all other high molecular weight PBPs (Δ5pbp). Each strain was grown to exponential phase, spotted on a 1% TY agar pad containing 1mM IPTG if inducing *mrcA*, allowed to grow for 16 hours, and imaged with phase microscopy. C. Working model of activation of ChvG-ChvI in *A. tumefaciens*. H^+^, free proton representing an acidic environment; β, β-lactam antibiotic.

We hypothesized that the ChvG-ChvI pathway may confer resistance to cell wall stress in other host-associated Rhizobiales. Indeed, depletion of PBP1a in *Sinorhizobium meliloti* causes cells to spread suggest activation of ChvG-ChvI (ExoS-ChvI) pathway (Fig 6B). Like *A. tumefaciens*, this response is specific to depletion of PBP1a and is not triggered by deletion of the 5 other high molecular weight PBPs. These results suggest that cell wall stress may be a well conserved trigger for activation of ChvG-ChvI pathways in the Rhizobiales. Notably, *C. crescentus* does not spread upon treatment with 80 μg/mL of cefsulodin (Fig S6C), despite recent findings that ChvGI likely confers resistance at this concentration [12]. While *S. meliloti* and *A. tumefaciens* encode succinoglycan biosynthesis operons, *C. crescentus* does not. These findings support the hypothesis that the ChvG-ChvI response to cell wall stress is conserved across Alphaproteobacterial species that have orthologs of ChvG-ChvI.

## DISCUSSION

Why is ChvG-ChvI TCS activated upon inhibition of polar PG synthesis in *A. tumefaciens*? PBP1a depletion results in a compositional shift in cell wall muropeptide composition and cell wall crosslinking [16]. Here we find that PBP1a depletion causes upregulation in transcription of LD-transpeptidases, endopeptidases, and soluble lytic transglycosylases, indicating cells are attempting to compensate for compromised cells walls. Remarkably, transcripts from these same genes are upregulated in the Δ*exoR* and pH 5.5 datasets, suggesting that remodeling of the cell wall is a part of the ChvG-ChvI regulon. While the role of cell wall remodeling during host-invasion is unclear it is possible that these modifications may be protective for the bacterium when host associated. The upregulation in transcription of genes encoding AopB and the Tol/Pal system may indicate that PBP1a-depleted cells are succumbing to osmotic pressure, a possible explanation for the increase in cell width in PBP1a-depleted and cefsulodin-treated cells (Fig 5B). Notably, this concept aligns well with recent findings in *Caulobacter crescentus* where it was shown that ChvGI activation was required for osmoregulation [35]. Together, these findings indicate a conserved role within the Alphaproteobacteria for the ChvG-ChvI TCS in sensing and responding to envelope stress. Possible signals may include the accumulation of cytoplasmic peptidoglycan precursors such as lipid II, increased levels of naked glycan strands in the cell wall, the production of peptidoglycan hydrolysis products, or more conventional stress responses due to osmotic sensitivity [36]. Future investigations will be necessary to determine if ChvG-ChvI proteins themselves directly sense and respond to cell wall stress or if other regulators which, presumably interact with the ChvG-ChvI are involved in this response.

If there is a conserved role in sensing cell wall stress, why would this pathway be required for host invasion within *Agrobacterium tumefaciens*? One explanation could be that during plant colonization, *A. tumefaciens* decreases cell wall biogenesis to form cells which are relatively persistent in order to evade host recognition and survive the harsh *in plantae* environment [37]. Alternatively, perhaps the integrity and composition of the cell wall is routinely monitored and used as a signal for the activation of pathways associated with lifestyle choice. Decreased activity of PBP1a may mimic one or more of the conditions *A. tumefaciens* and *S. meliloti* encounter during host-invasion, leading to the activation of ChvG-ChvI. Alternatively, perhaps the absence of PBP1a activity leads to a destabilization of the polar growth complex leading to decreased cell envelope integrity. It will be of interest to determine if other components of the polar growth complex such as GPR [38], RgsP [39], or PopZ [40,41] have increased sensitivity to β-lactam antibiotics and induce surface spreading. This possibility is in agreement with the observation in *C. crescentus* that resistance to cell wall targeting antibiotics is dependent on factors such as TipN that maintain the integrity of the cell envelope [42].

The overproduction of succinoglycan provides cells with passive protection against several stresses *A. tumefaciens* may encounter during host invasion including detergents, salt, acidity, heat, antimicrobial peptides, and reactive oxygen species [33-36]. Production of succinoglycan may also help protect against cell-wall-synthesis targeting antibiotics produced by competing bacteria and fungi in the soil. However, succinoglycan production is taxonomically constrained within plant-host-associated bacteria, indicating that there are other conserved mechanisms regulated by ChvG-ChvI in resistance to these antibiotics. The surface spreading phenomenon that we connected to overproduction of succinoglycan and that has been previously described in *S. meliloti* [30], may be involved in cell dispersal during host invasion.

The increased sensitivity to mecillinam, vancomycin, cefsulodin, and moenomycin in ChvGI mutants suggests that this pathway may confer resistance to antibiotics inhibiting cell wall synthesis [35]. Our observation that the depletion of *A. tumefaciens* RgsM mutant does not induce surface spreading further hints that increased cell wall hydrolysis may be responsible for activation of the ChvG-ChvI pathway. While our findings suggest a conserved signal in ChvG-ChvI activation, further studies will need to be conducted to identify the signal(s), determine if they are species-specific, and explore the conditions which lead to signal production.

Longer depletions of PBP1a may induce a more general stress response. While previous reports propose *A. tumefaciens* to have stress-specific responses instead of a single general response [43], the similarities between the transcriptional changes at the 16-hour time point and the general stress responses of other bacteria are striking [44]. For example, the general stress response of *Escherichia coli*, regulated by the ECF family sigma factor σ^E^, upregulates transcription of genes encoding ABC transporters associated with uptake of amino acids, peptides, phosphate sources, and polyamines [45]. Indeed, the 16-hour time point shows differential expression of over 30 ABC transporters. However, previous studies found no common response to stresses including osmotic stress, heat shock, and low pH in *A. tumefaciens* [43]. While there are several ECF family sigma factors encoded in the *A. tumefaciens* genome, it is unclear what, if any, role they may be playing in the transcriptional changes exhibited under long depletions of PBP1a.

Together our findings support a model that ChvG-ChvI can be acid induced by dissociation and proteolysis of ExoR in a Rhizobiales specific response (Fig 6C, left) and through a more broadly conserved cell envelope stress response (Fig 6C, middle). The role of ExoR regulation is well established for the acid-induction of ChvG-ChvI but the lack of ExoR conservation suggests that the cell envelope stress response may be ExoR-independent. Finally, longer periods of PBP1a depletion results in a more general stress response, affecting transcription of genes associated with nutrient uptake and metabolism (Fig 6C, right). Overall, the data presented here are in agreement with recent works in *Caulobacter* [35,42] which suggest that ChvGI activation is important in oligotrophic free-living bacteria as a cell envelope or osmotic stress response. Remarkably, it seems that the ChvG-ChvI pathway has a dual purpose in protecting the bacterium and invading its host in *A. tumefaciens*, and other host-associated Rhizobiales.

## MATERIALS AND METHODS

### Bacterial strains, plasmids, and growth conditions

A list of all bacterial strains and plasmids used in this study is provided in Table S2. *Agrobacterium tumefaciens* C58 and derived strains were grown in ATGN minimal media [46] without exogenous iron at 28°C with shaking. When appropriate, kanamycin (KAN) was used at the working concentration of 300 μg/ml. When indicated, isopropyl β-D-1-thio-galactopyranoside (IPTG) was used as an inducer at a concentration of 1 mM, and cumate. *Sinorhizobium meliloti* stains were grown in TY medium [47] at 28°C. When appropriate, KAN was used at the working concentration of 200 μg/ml, gentamycin (GM) was used at 20 μg/ml, and IPTG was used at a concentration of 500 μg/ml. *C. crescentus* strains were grown in PYE medium [48] at 28°C. *E. coli* DH5α and S17-1 λ pir were grown in Luria-Bertani medium at 37°C and when appropriate 50 μg/ml or 30 μg/ml of KM were added, respectively.

### Construction of deletion/depletion plasmids and strains

A list of all primers and synthetic DNAs used in this study is provided in Table S3. Vectors for gene deletion by allelic exchange were constructed using recommended methods for *A. tumefaciens* [49]. Briefly, 500-bp fragments upstream and 500 bp downstream of the target gene were amplified using primer pairs P1/P2 and P3/4 respectively. Amplicons were spliced together by SOEing using primer pair P1/P4. The amplicon was digested and ligated into pNTPS139. The deletion plasmids were introduced into *A. tumefaciens* by mating using an *E. coli* S17 conjugation strain to create KM resistant, sucrose sensitive primary integrants. Primary integrants were grown overnight in media with no selection. Secondary recombinants were screened by patching for sucrose resistance and KM sensitivity. Colony PCR with primers P5/P6 for the respective gene target was used to confirm deletion. PCR products from P5/P6 primer sets were sequenced to further confirm deletions.

### Phase and fluorescence microscopy

A small volume (∼1 μl) of cells in exponential phase (OD600 = 0.2 - 0.4) was applied to a 1% ATGN agarose pad as described previously [50]. DIC, Phase contrast and epifluorescence microscopy were performed with an inverted Nikon Eclipse TiE and a QImaging Rolera em-c2 123 1K EMCCD camera with Nikon Elements Imaging Software. For time-lapse microscopy, images were collected every ten minutes, unless otherwise stated. For calcofluor agar pad assays, calcofluor was added to agarose pads at a concentration of 25 μg/mL and exposed to DAPI filter for 50ms. When appropriate agar pads were supplemented with 1mM IPTG. For quantitative image analysis, live cells were imaged using phase-contrast microscopy, and cell length and width distributions of the indicated number of cells per strain were determined as measured using MicrobeJ software [51]. T-tests were performed using the compare_means() function of the ggpubr R library.

### RNA isolation, sequencing and analysis

Four cultures each of WT, WT + IPTG and 12 cultures of + PBP1a depletion cells were grown overnight in 2 ml of ATGN minimal media at 28°C with shaking; the + PBP1a strains and WT + IPTG strains were supplemented with 1mM IPTG. Cells were then pelleted by centrifugation at 7000 x g for 5 minutes. Cell pellets were washed three times with ATGN by centrifugation and resuspension to remove IPTG. After the final wash the cell pellets from WT, WT + IPTG, and four of the 12 + PBP1a strains were resuspended to an OD600 of 0.05 in 6mL ATGN, or ATGN with 1mM IPTG. The other eight + PBP1a strains were resuspended to an OD600 of 0.05 in 6 ml ATGN without IPTG. This resulted in 4 replicate cultures each of WT, WT +IPTG, + PBP1a, - PBP1a_6hr and - PBP1a_16hr. Growth of the cultures was monitored and supplemented with fresh medium as needed so that the OD600 never went over 0.3. RNA was isolated from the -PBP1a_6hr strains after 6 hours of growth, and RNA was isolated from all other strains after 16 hours of growth. To prepare samples, a culture volume equivalent to 6 ml at an optical density at 600 nm (OD600) of 0.2-0.3 was pelleted by centrifugation at 7000 x g for 5 minutes and pellets were resuspended in 1mL of ATGN media and incubated with 2 mL of RNAProtect reagent (QIAgen) for 15 min at room temperature. Cells were lysed with 10 mg lysozyme, and RNA was extracted using the QIAgen RNEasy kit.

DNA libraries for sequencing were constructed following the manufacturer’s protocol with reagents supplied in Illumina’s TruSeq mRNA stranded sample preparation kit. The sample concentration was determined by Qubit flourometer (Invitrogen) using the Qubit HS RNA assay kit, and the RNA integrity was checked using the Fragment Analyzer automated electrophoresis system. Briefly, the poly-A containing mRNA is purified from total RNA, RNA is fragmented, double-stranded cDNA is generated from fragmented RNA, and the index containing adapters are ligated to the ends. The amplified cDNA constructs were purified by addition of Axyprep Mag PCR Clean-up beads. The final construct of each purified library was evaluated using the Fragment Analyzer automated electrophoresis system, quantified with the Qubit flourometer using the Qubit HS dsDNA assay kit, and diluted according to Illumina’s standard sequencing protocol for sequencing on the NextSeq 500.

For all samples, when adapter sequence was detected, it was removed using cutadapt (0.16) [52]. All samples were purged of reads that mapped to transcripts for rRNA genes using bowtie2 (2.3.4.3) [53]. The reads were then mapped to the *A. fabrum* str. C58 genome using STAR (version 2.5.4b) [54], which also produces the number of read counts per gene. The index files used by STAR were derived from the files *Agrobacterium*_fabrum_str_c58.ASM9202v1.dna.toplevel.fa and *Agrobacterium*_fabrum_str_c58.ASM9202v1.40.gtf, both of which are part of Ensembl release 40 (http://bacteria.ensembl.org/index.html). Pairwise comparisons were performed to test for differential expression of genes using the Bioconductor package DESeq2 [55]. Gene annotations were collected from the annotations included with the file of cDNAs also at Ensembl *Agrobacterium*_fabrum_str_c58.ASM9202v1.cdna.all.fa.gz.

### COG functional annotation

Amino acid sequences for all proteins in *A. tumefaciens* were downloaded in a single FASTA file from GenBank and uploaded to EGGNOG-MAPPER [56,57]. COG terms were outputted, and Python code was written to pull out transcripts from 16-hour depletion of PBP1a with L2FC > 2.0 or < −2.0. Some transcripts had multiple COG annotations and were therefore replicated for visualization according to the number of annotations it had.

### Comparative transcriptomics

Transcripts and L2FC values from each dataset were opened in Python code written to screen for and exclude any genes that were not present in both datasets. Statistics and visualization was done in R. Spearman correlation statistical test was run on the L2FC of the 150 most differentially expressed genes in the PBP1a depletion and their corresponding L2FC values in the comparison dataset.

### Cell viability assays

For cell viability spot assays, cultures were grown overnight and diluted to an OD_600_ = 0.05 and serially diluted in ATGN and spotted on ATGN agar plates containing antibiotics as indicated. Four microliters of each dilution was spotted and plates were incubated at 28°C for 48 h before imaging.

### Disk diffusion assays

Wild-type, Δ*chvG*, and Δ*chvI* cells were overnight and then knocked down to an OD_600_ of 1.0. Cells were then lawned on ATGN minimal media. Sterile paper disks either soaked in concentrations of each antibiotic or not (blank controls) were applied to the plate. Each plate was grown for ∼48 hours at 28°C before being imaged. Zone of inhibition diameters were measured from each image using ImageJ software.

### Phylogenetics and structure prediction

A seed of 22 amino acid sequences containing the annotated ChvG sensor domain (PF13755) were initially downloaded from Pfam [58]. Each was blasted against its corresponding proteome to retrieve the full protein sequence [59]. Additional sequences of relevant bacteria such as *S. meliloti, Brucella melitensis*, and *C. crescentus*, were added by blasting the amino acid sequence from *A. tumefaciens* ChvG (Atu0033) against each organism’s proteome. All sequences were aligned using MUSCLE and trimmed in Jalview according to Uniprot predicted periplasmic region of Atu0033 [60–62]. A maximum parsimony phylogenetic tree of these sequences was generated using MEGA-X [63].

Each trimmed sequence underwent one-to-one threading in Phyre2 with the complete structure of Atu0033 predicted by AlphaFold as a template [64,65]. Local alignment and a secondary structure weight of 0.1 was used. Structural analysis and structure alignment was done in ChimeraX [66].

The amino acid sequence of *A. tumefaciens* ExoR (Atu1715) was blasted against each organism’s proteome and max score values of top hits were recorded. Max score values under 50 were deemed too different and were therefore not considered an ExoR ortholog. Additionally, sequences of each top hit were blasted against the proteome of *A. tumefaciens*. If the top hit was not ExoR, it was also not considered an ExoR ortholog in this analysis.

## DATA AVAILABILITY

Raw RNA-seq read files (.fastq) and complete list of differentially expressed genes for each comparison are publicly available through the NCBI Gene Expression Omnibus (GEO) under the accession <u>GSE173921</u>.

## ACKNOWLEDGEMENTS

We thank Elizaveta Krol and Anke Becker for providing *Sinorhizobium meliloti* strains and micrographs and Erh-Min Lai for providing antibodies used in this work. We thank Amelia Randich, Alex Quintero, Regis Hallez and members of the Brown lab for critical feedback on this manuscript. This work was supported by the National Science Foundation, IOS1557806, to PJBB. MAW and JMB were supported by the Life Sciences Fellowship at the University of Missouri and AKM was supported as an IMSD/MARC fellow with support from the National Institutes of Health, T34 GM136493.

The molecular graphics and analyses were performed with UCSF ChimeraX, developed by the Resource for Biocomputing, Visualization, and Informatics at the University of California, San Francisco, with support from National Institutes of Health R01-GM129325 and the Office of Cyber Infrastructure and Computational Biology, National Institute of Allergy and Infectious Diseases.

## SUPPORTING INFORMATION

**Movie S1:** Growth and morphological changes during 8 hours of PBP1a depletion. Cells were washed to remove inducer and spotted immediately on an ATGN pad. Images were acquired every ten minutes and movie is played at 16 frames per second for a total of 48 frames.

**Movie S2:** Growth and division of D*chvI* during PBP1a depletion. Cells were washed to remove inducer and spotted immediately on an ATGN pad. Images were acquired every five minutes and movie is played at 40 frames per second for a total of 200 frames.

**Fig S1. Pelleting of PBP1a-depleted and repleted cells**. Conical tubes show turbidity after pelleting cells grown for 16 hours in PBP1a replete (+PBP1a) or depleted (-PBP1a) conditions. Cells were centrifuged at 1690 x g (3000 rpm in TX-400 rotor in a Sorvall Legend X1R centrifuge) for 10 minutes. Supernatants were spotted on a 1.25% ATGN agarose pad. x = average number of cells from 10 fields of view.

**Fig S2. Analysis of the control transcriptomes by RNA-seq**. A. Plots comparing Log2Fold Change of the WT 6-hour transcriptome to that of the WT +ITPG 6-hour transcriptome. Gray dots represent a single transcript, and the dotted lines represent +/-2.0 Log2Fold Change threshold. Plots are delimited by chromosomes and mega plasmids. B. Plots comparing Log2Fold Change of the WT +IPTG transcriptome to that of the PBP1a depletion strain with ITPG present to drive PBP1a expression. Comparisons shown are of the 6-hour transcriptomes. Gray dots represent a single transcript, and the dotted lines represent +/-2.0 Log2Fold Change threshold. Plots are delimited by chromosomes and mega plasmids.

**Fig S3. Transcriptional changes of TCS regulators and kinases during PBP1a depletion**. The fold change in expression level of TCS regulators and kinases are shown following 6 hours (gray) and 16 hours (black) of PBP1a depletion. The *virAG* and *chvGI* TCS pairs are labeled.

**Fig S4. The response to the depletion of PBP1a mimics transcriptional changes associated with host invasion**. A. Correlation scatterplots depicting relationships between the log2fold-change (L2FC) values in the 16-hour PBP1a depletion and transcriptomic data sets taken under simulated virulence-inducing conditions (AS) and under simulated host-invading conditions (Δ*exoR*). Each point represents a unique transcript. AS, acetosyrinogone; Rho, Spearman correlation coefficient. B. Correlation scatterplots comparing L2FC values of transcripts in the pH 5.5 microarray, a condition known to induce the *chvG-chvI* regulon, to either the 6-hour (red) or 16-hour (blue) PBP1a depletion. Rho, Spearman correlation coefficient.

**Fig S5. Western blot of proteins expressed from the two type VI secretion system operons in ΔT6SSpro strains**. Top panel, diagram of the two operons encoding elements of Type VI Secretion in *A. tumefaciens*. T6SSpro labels the intergenic gap that is deleted in ΔT6SSpro strains. Middle panel, western blots using anti-Hcp and anti-TssB in each of the indicated strains. Protein sizes (kDa) are shown on the right. Bottom panel, Coomassie stained gel showing total protein from each strain.

**Fig S6. Impact of decreased PG synthesis on *A. tumefaciens* and *C. crescentus* microcolony formation**. A. Micrographs of PBP1a depletion Δ*chvI* with (+PBP1A) or without (-PBP1A) IPTG. Cells was grown to exponential phase in ATGN media containing IPTG, spotted on an ATGN agar pad with or without IPTG, allowed to grow for 16 hours, and imaged by DIC microscopy. B. Micrographs of WT *A. tumefaciens* cells growth with or without moenomycin. Cells was grown to exponential phase in ATGN media, spotted on an ATGN agar pad with or without moenomycin, allowed to grow for 16 hours, and imaged by DIC microscopy. C. Micrographs of WT *C. crescentus* cells growth with or without cefsulodin. Cells were grown to exponential phase in PYE media, spotted on a PYE agar pad with or without cefsulodin, allowed to grow for 16 hours, and imaged by DIC microscopy. All scale bars depict 2μm.

**Fig S7. Alignment of periplasmic regions of ChvG orthologs**. Partial MUSCLE alignment of ChvG ortholog periplasmic domains. Highlighted columns represent strong conservation across aligned sequences. Atu0033 (ChvG of *A. tumefaciens)* is the reference sequence for this analysis. L1 and L2 correspond to two conserved structural loops. Conservation, quality, and consensus scores for each site are represented as bar graphs under the alignment. Shading indicates order of the bacterium containing the ChvG ortholog: Orange, Rhizobiales; Purple, Rhodobacterales; Green, Caulobacterales; Gold, Rhodospirales.

**Fig S8. Structure predictions for the periplasmic regions of ChvG orthologs**. Phyre2 structural predictions for each organism displayed in Figure 6A of this work. Genus and species names as well as locus tags for each ChvG ortholog are provided. Range of numbers following the back slash are the amino acid sites used in structure prediction. Colors indicates order of the bacterium containing the ChvG ortholog: Orange, Rhizobiales; Purple, Rhodobacterales; Green, Caulobacterales; Gold, Rhodospirales.

**Table S1. Selected differentially expressed genes**

**Table S2. Bacterial strains and plasmids**.

**Table S3. Synthesized DNA primers.**

**Supplementary Methods**

**Supplementary References**

## Notes

### Competing Interest Statement

The authors have declared no competing interest.

### Summary of Updates

Addition of supplemental material

