## Supplemental Material for "Activation of ChvG-ChvI regulon by cell wall stress confers resistance to β-lactam antibiotics and initiates surface spreading in *Agrobacterium tumefaciens*"

**Title:**

**Running title:** Cell wall stress activates the ChvG-ChvI pathway in *A. tumefaciens*

*denotes equal contributions

**^#^Corresponding author:**

(573) 884-0384

**Present address:**

^$^Boulevard Brewery, 2501 Southwest Blvd, Kansas City, MO 64108

**SUPPLEMENTARY TEXT**

1) **Supplementary Movie Legends**
2) **Supplementary Figures and Legends**
3) **Supplementary Tables**
   a. **Table S1. Selected differentially expressed genes**
   b. **Table S2. Bacterial strains and plasmids**
   c. **Table S3. Synthesized DNA primers**
4) **Supplementary Methods**
5) **Supplementary References**

**1) Supplementary Movie Legends**

**Movie S1.** Growth and morphological changes during 8 hours of PBP1a depletion. Cells were washed to remove inducer and spotted immediately on an ATGN pad. Images were acquired every ten minutes and movie is played at 16 frames per second for a total of 48 frames.

**Movie S2.** Growth and division of Δ*chvI* during PBP1a depletion. Cells were washed to remove inducer and spotted immediately on an ATGN pad. Images were acquired every five minutes and movie is played at 40 frames per second for a total of 200 frames.

**2) Supplementary Figures and Legends**

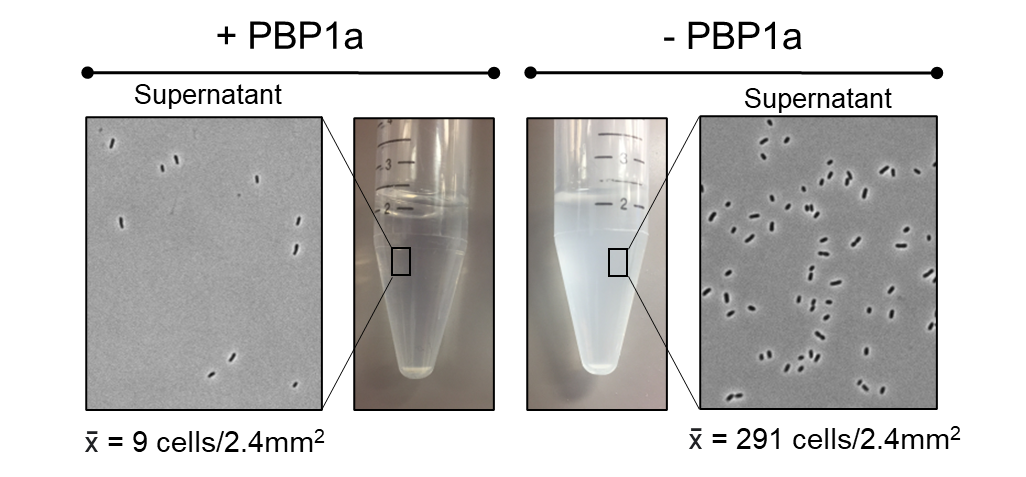

**Fig S1.** **Pelleting of PBP1a-depleted and repleted cells.** Conical tubes show turbidity after pelleting cells grown for 16 hours in PBP1a replete (+PBP1a) or depleted (-PBP1a) conditions. Cells were centrifuged at 1690 x g (3000 rpm in TX-400 rotor in a Sorvall Legend X1R centrifuge) for 10 minutes. Supernatants were spotted on a 1.25% ATGN agarose pad. x̄ = average number of cells from 10 fields of view.

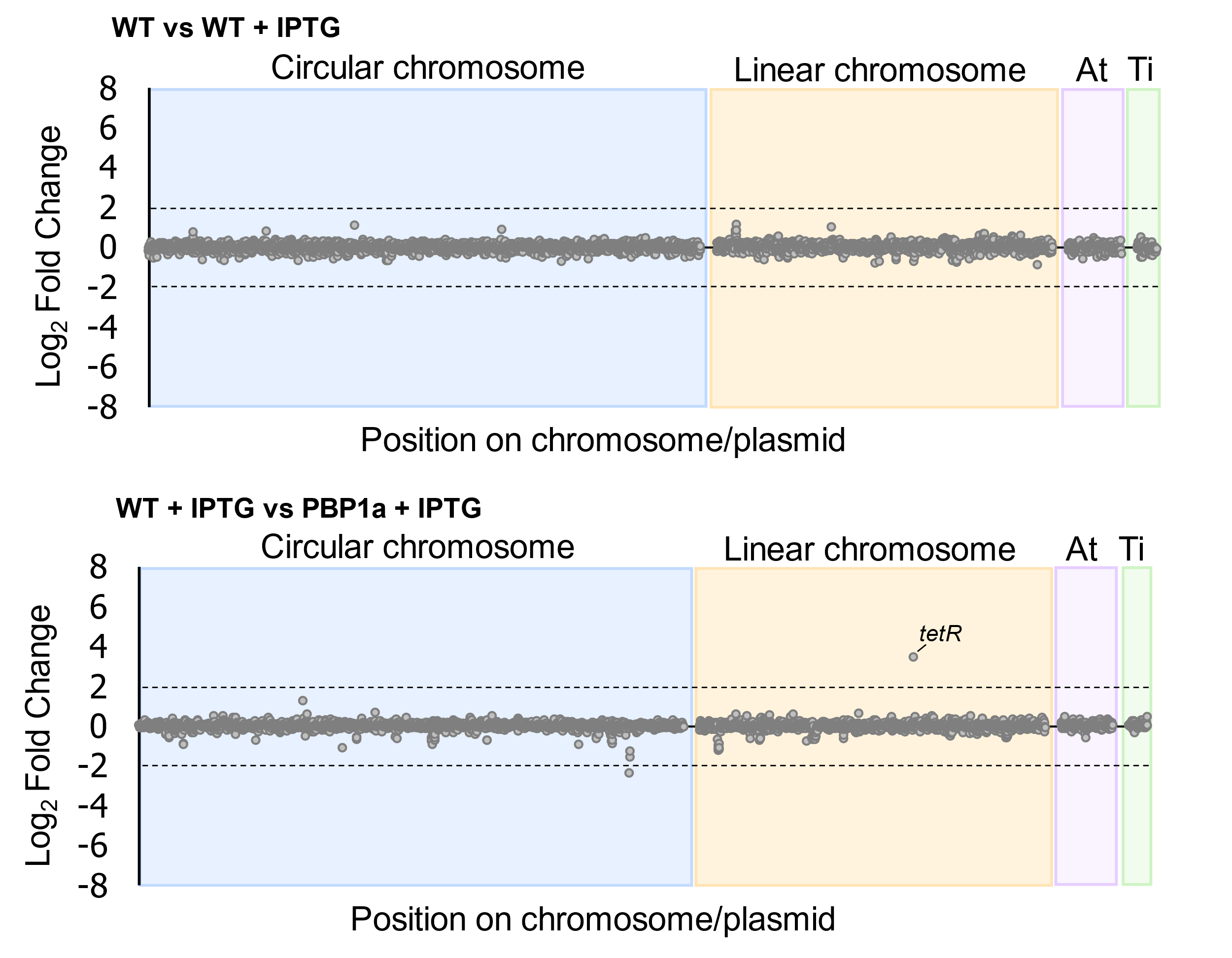

**B**

**A**

**Fig S2.** **Analysis of the control transcriptomes by RNA-seq.** A. Plots comparing Log2Fold Change of the WT 6-hour transcriptome to that of the WT +ITPG 6-hour transcriptome. Gray dots represent a single transcript, and the dotted lines represent +/- 2.0 Log2Fold Change threshold. Plots are delimited by chromosomes and mega plasmids. B. Plots comparing Log2Fold Change of the WT +IPTG transcriptome to that of the PBP1a depletion strain with ITPG present to drive PBP1a expression. Comparisons shown are of the 6-hour transcriptomes. Gray dots represent a single transcript, and the dotted lines represent +/- 2.0 Log2Fold Change threshold. Plots are delimited by chromosomes and mega plasmids.

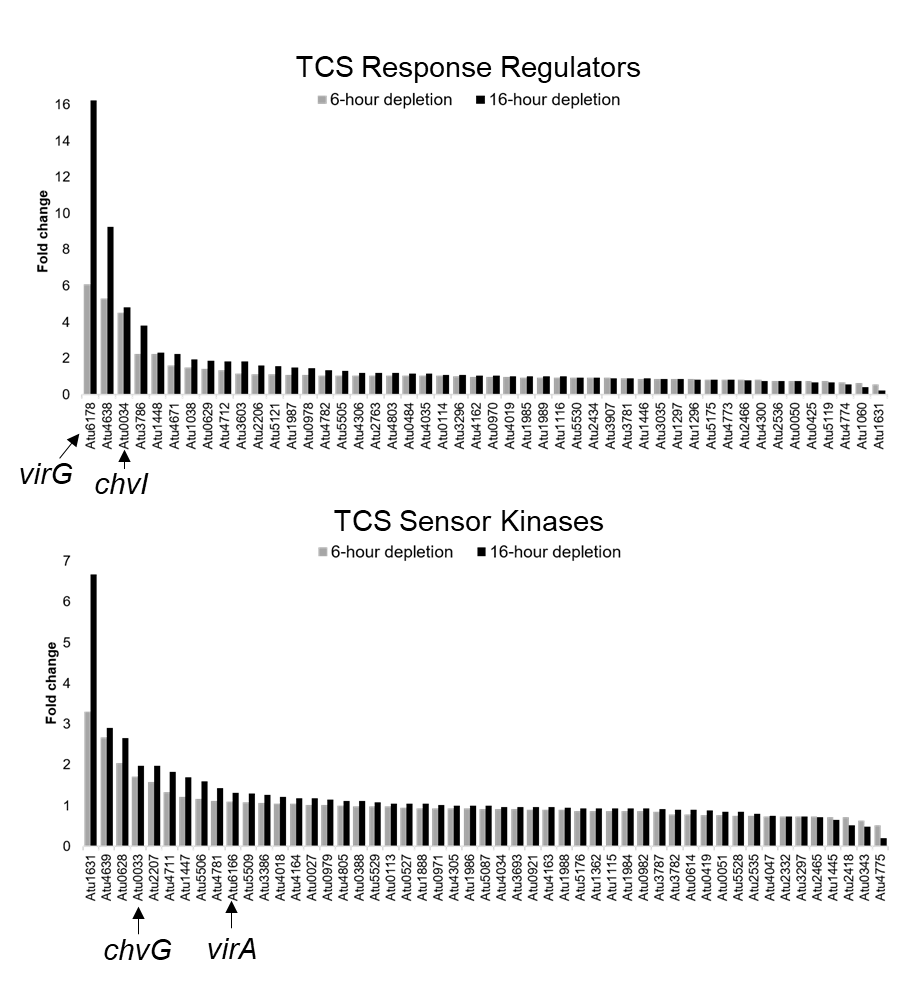

**Fig S3. Transcriptional changes of TCS regulators and kinases during PBP1a depletion.** The fold change in expression level of TCS regulators and kinases are shown following 6 hours (gray) and 16 hours (black) of PBP1a depletion. The *virAG* and *chvGI* TCS pairs are labeled.

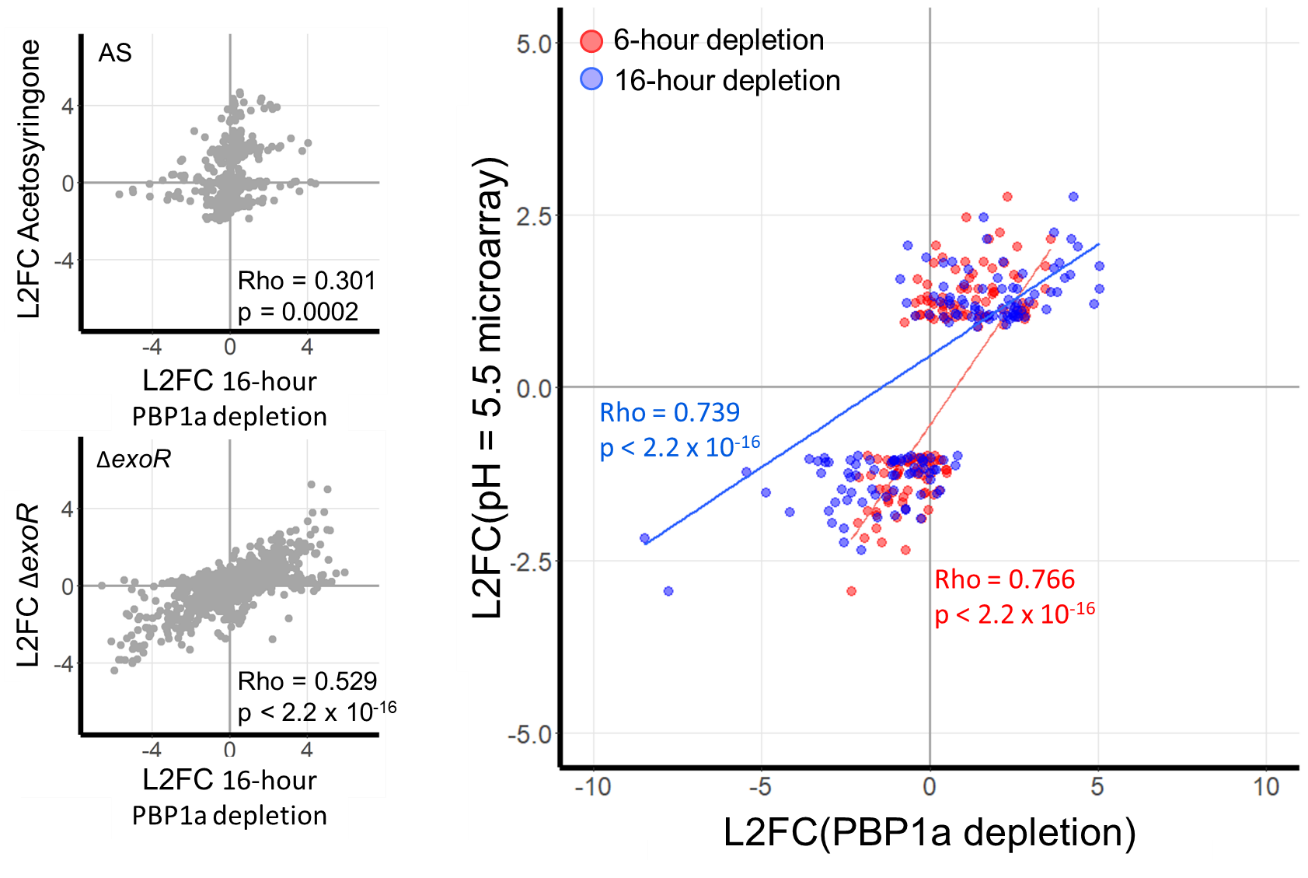

**B**

**A**

**Fig S4. The response to the depletion of PBP1a mimics transcriptional changes associated with host invasion.** A. Correlation scatterplots depicting relationships between the log2fold-change (L2FC) values in the 16-hour PBP1a depletion and transcriptomic data sets taken under simulated virulence-inducing conditions (AS) and under simulated host-invading conditions (Δ*exoR*). Each point represents a unique transcript. AS, acetosyrinogone; Rho, Spearman correlation coefficient. B. Correlation scatterplots comparing L2FC values of transcripts in the pH 5.5 microarray, a condition known to induce the *chvG-chvI* regulon, to either the 6-hour (red) or 16-hour (blue) PBP1a depletion. Rho, Spearman correlation coefficient.

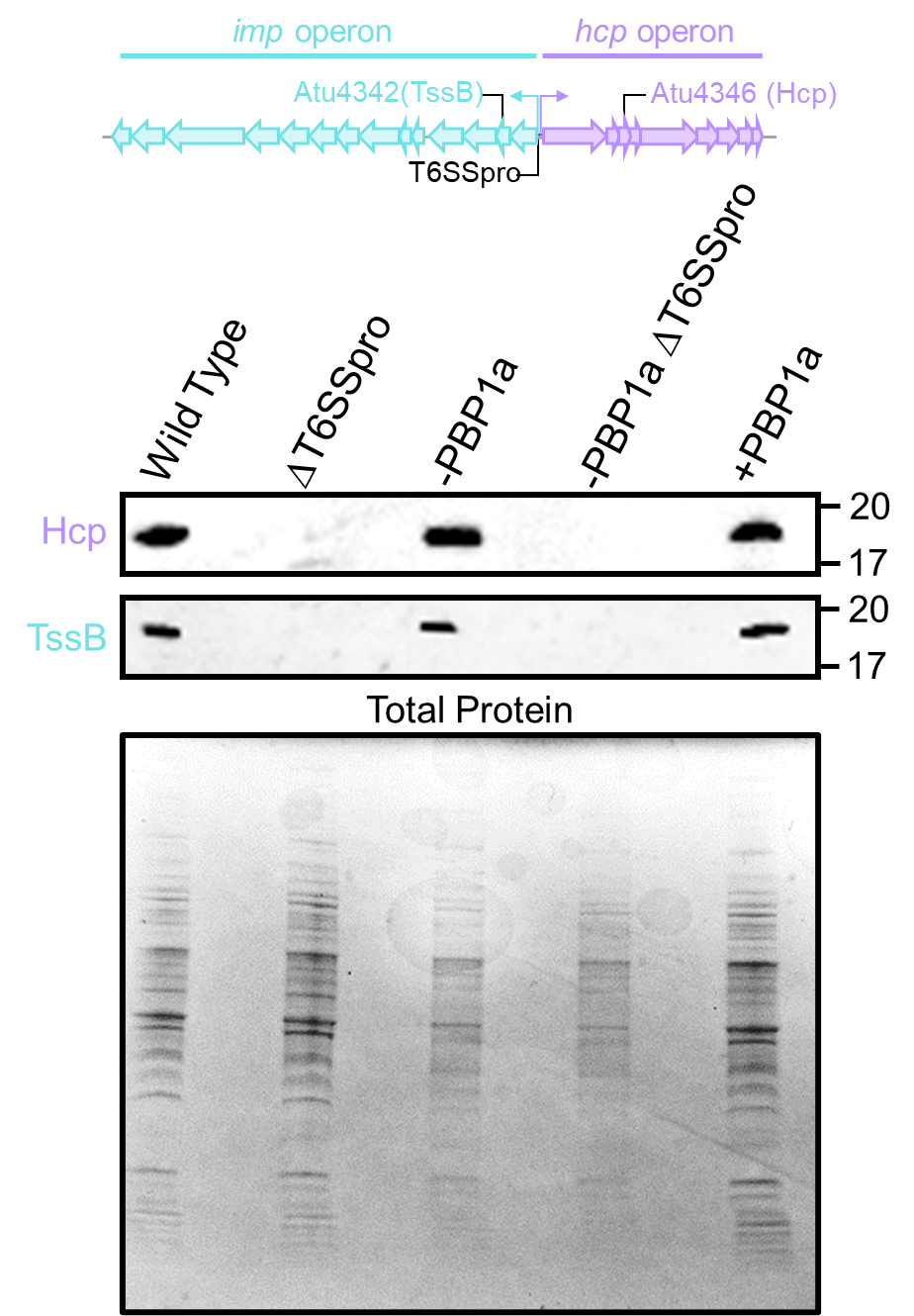

**Fig S5.** **Western blot of proteins expressed from the two type VI secretion system operons in ΔT6SSpro strains.** Top panel, diagram of the two operons encoding elements of Type VI Secretion in *A. tumefaciens.* T6SSpro labels the intergenic gap that is deleted in ΔT6SSpro strains. Middle panel, western blots using anti-Hcp and anti-TssB in each of the indicated strains. Protein sizes (kDa) are shown on the right. Bottom panel, Coomassie stained gel showing total protein from each strain.

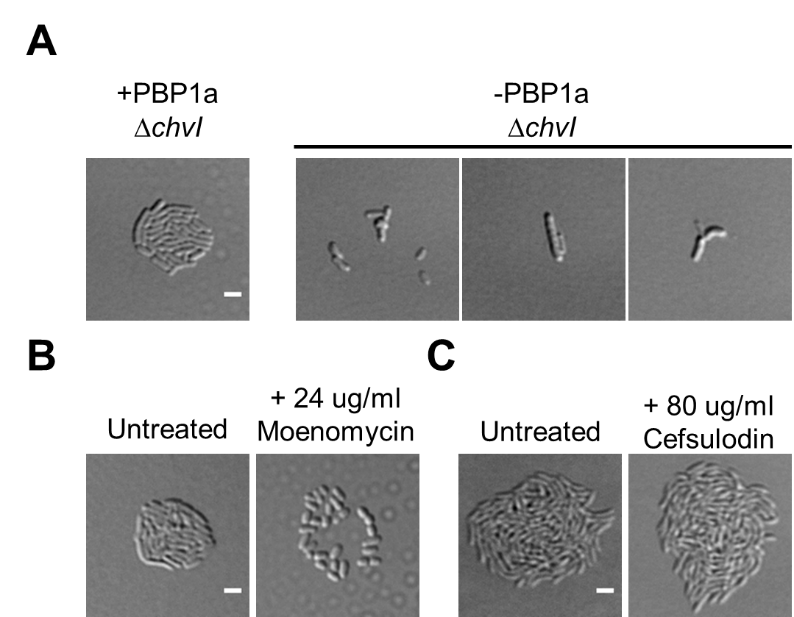

**Fig S6. Impact of decreased PG synthesis on *A. tumefaciens* and *C. crescentus* microcolony formation.** A. Micrographs of PBP1a depletion Δ*chvI* with (+PBP1A) or without (-PBP1A) IPTG. Cells was grown to exponential phase in ATGN media containing IPTG, spotted on an ATGN agar pad with or without IPTG, allowed to grow for 16 hours, and imaged by DIC microscopy. B. Micrographs of WT *A. tumefaciens* cells growth with or without moenomycin. Cells was grown to exponential phase in ATGN media, spotted on an ATGN agar pad with or without moenomycin, allowed to grow for 16 hours, and imaged by DIC microscopy. C. Micrographs of WT *C. crescentus* cells growth with or without cefsulodin. Cells were grown to exponential phase in PYE media, spotted on a PYE agar pad with or without cefsulodin, allowed to grow for 16 hours, and imaged by DIC microscopy. All scale bars depict 2μm.

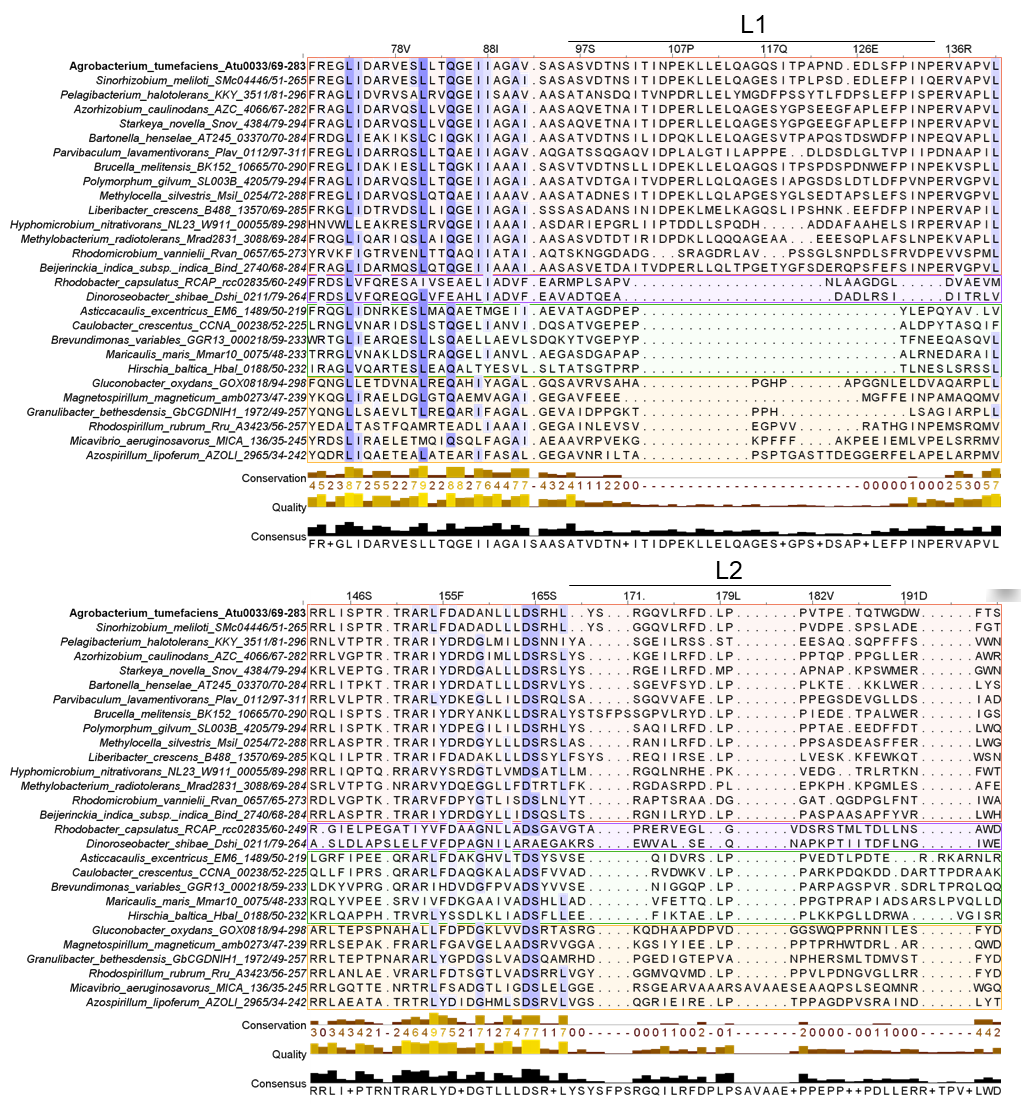
**Fig S7. Alignment of periplasmic regions of ChvG orthologs.** Partial MUSCLE alignment of ChvG ortholog periplasmic domains. Highlighted columns represent strong conservation across aligned sequences. Atu0033 (ChvG of *A. tumefaciens)* is the reference sequence for this analysis*.* L1 and L2 correspond to two conserved structural loops. Conservation, quality, and consensus scores for each site are represented as bar graphs under the alignment. Shading indicates order of the bacterium containing the ChvG ortholog: Orange, Rhizobiales; Purple, Rhodobacterales; Green, Caulobacterales; Gold, Rhodospirales.

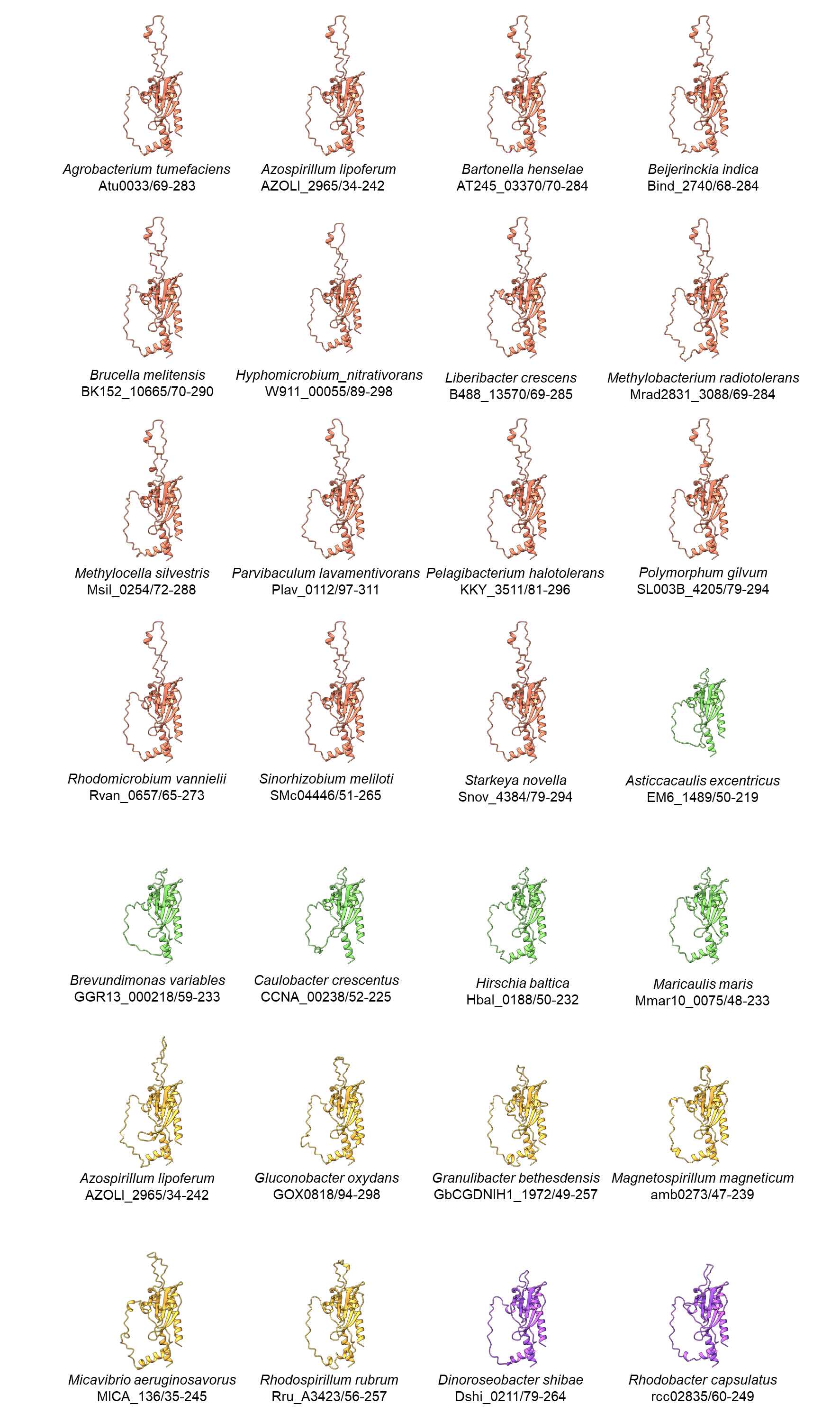

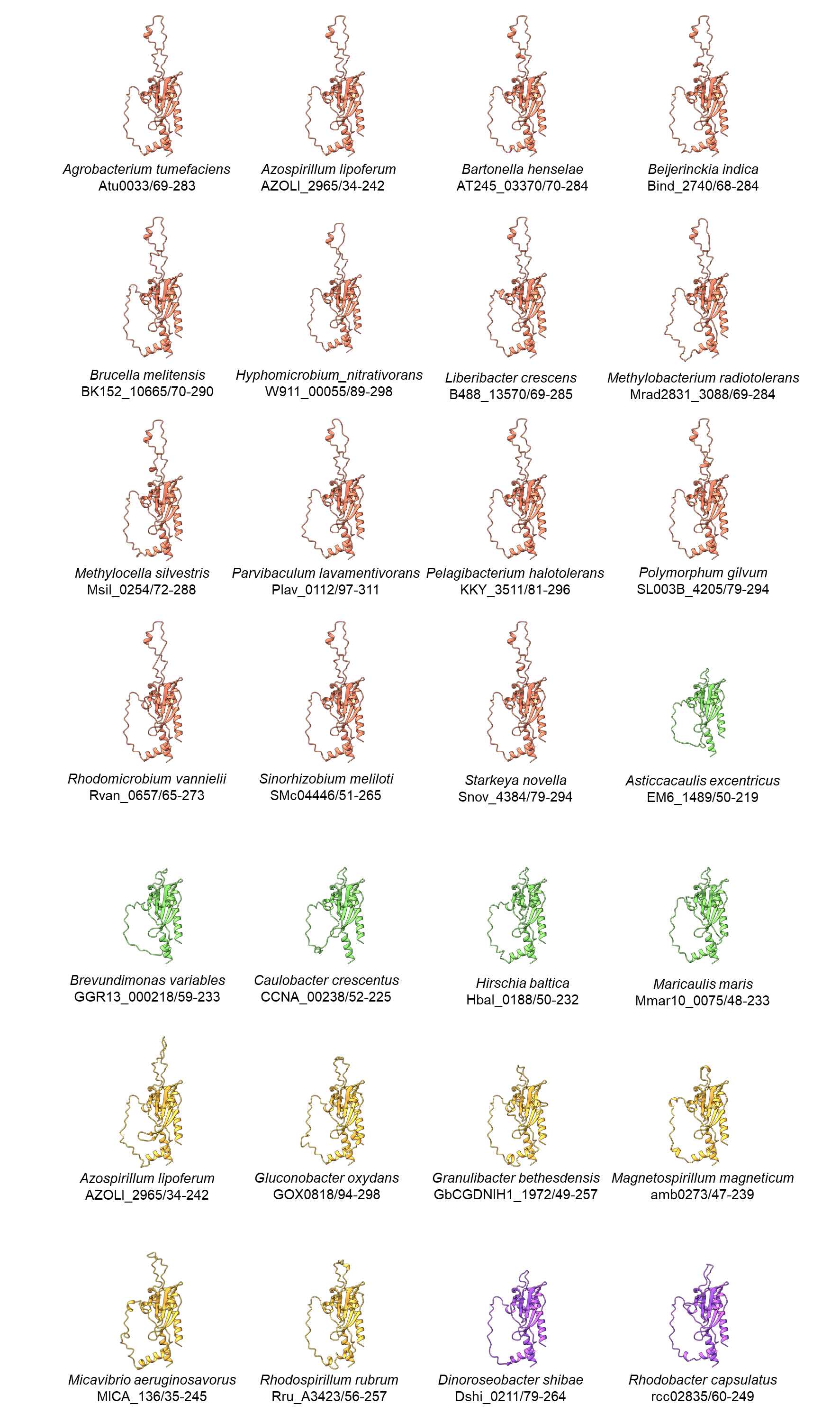

**Fig S8. Structure predictions for the periplasmic regions of ChvG orthologs.**

Phyre2 structural predictions for each organism displayed in Figure 6A of this work. Genus and species names as well as locus tags for each ChvG ortholog are provided. Range of numbers following the back slash are the amino acid sites used in structure prediction. Colors indicates order of the bacterium containing the ChvG ortholog: Orange, Rhizobiales; Purple, Rhodobacterales; Green, Caulobacterales; Gold, Rhodospirales.

**3) Supplementary Tables**

**Table S1. Selected differentially expressed genes**

| **Gene Name** | **GeneID** | **Description** | **log_2_ FC** | **p value** |
| --- | --- | --- | --- | --- |
| **Cell wall synthesis and remodeling** | | | | |
| *pbp1a* | Atu1341 | class A penicillin-binding protein 1a | -7.20 | 0 |
| *pbp1b1* | Atu0103 | class A penicillin-binding protein 1b | 0.78 | 1.19E-45 |
| *pbp1b2* | Atu0931 | class A penicillin-binding protein 1b | 0.55 | 5.5E-14 |
| *pbp1c* | Atu3694 | class A penicillin-binding protein 1c | 0.12 | 0.10308 |
| *mtgA* | Atu2720 | monofunctional glycosyltransferase | -0.19 | 0.058441 |
| *pbp3a* | Atu2100 | class B penicillin-binding protein 3a | 0.71 | 2.86E-13 |
| *pbp3b* | Atu1064 | class B penicillin-binding protein 3b | -0.14 | 0.086185 |
| *ftsW* | Atu2095 | SEDS protein | 0.94 | 2.7E-23 |
| *rgsM* | Atu4178 | DD-endopeptidase | 1.72 | 5.4E-132 |
| *-* | Atu1832 | DD-endopeptidase | 0.41 | 0.01114 |
| *mepA* | Atu0186 | DD/LD-endopeptidase | 1.93 | 1.89E-77 |
| *-* | Atu2133 | LD-transpeptidase | 1.84 | 5.47E-85 |
| *-* | Atu3332 | LD-transpeptidase | 1.91 | 2.29E-114 |
| *-* | Atu0844 | LD-transpeptidase | -4.95 | 4.51E-90 |
| *-* | Atu2336 | LD-transpeptidase | -1.99 | 5.57E-51 |
| *-* | Atu0290 | soluble lytic transglycosylase | 2.46 | 4.7E-16 |
| *-* | Atu0572 | soluble lytic transglycosylase | -4.65 | 3.7E-125 |
| - | Atu2112 | soluble lytic transglycosylase | 5.17 | 1.1E-104 |
| - | Atu1221 | NLpC/p60 superfamily | 4.21 | 9.04E-21 |
| - | Atu0933 | beta-lactamase class D | 2.37 | 2.22E-54 |
| **Cell envelope homeostasis** | | | | |
| *palA* | Atu3713 | omp16 protein | 1.64 | 2.58E-51 |
| *tolB* | Atu3714 | tolB protein | 2.26 | 1.55E-116 |
| *tolA* | Atu3715 | conserved hypothetical protein | 2.06 | 1.98E-293 |
| *tolR* | Atu3716 | tolR protein | 2.06 | 3.97E-143 |
| *tolQ* | Atu3717 | tolQ Protein | 1.74 | 7.55E-34 |
| *ropB* | Atu1131 | OmpA-like outer membrane protein | 5.04 | 1.4E-149 |
| *-* | Atu1877 | OmpA-like outer membrane protein | 3.30 | 3.5E-159 |
| *-* | Atu1155 | periplasmic sensor creD | 4.38 | 0 |
| *-* | Atu2760 | lipoprotein transporter | 2.57 | 3.4E-108 |
| **Signaling** | | | | |
| *virA* | Atu6166 | two component sensor kinase | 0.52 | 1.56E-13 |
| *virG* | Atu6178 | two component response regulator | 4.02 | 0 |
|  | Atu4639 | two component sensor kinase | 2.74 | 2.9E-99 |
|  | Atu4638 | two component response regulator | 3.21 | 2.32E-68 |
| *rem* | Atu0573 | OmpR-type transcriptional regulator | -2.58 | 4.53E-89 |
| *visN* | Atu0524 | LuxR-type transcriptional regulator | -0.17 | 0.034462 |
| *visR* | Atu0525 | LuxR-type transcriptional regulator | 0.16 | 0.114024 |
| *exoR* | Atu1715 | eps production negative regulator | 2.20 | 1.64E-37 |
| *chvG* | Atu0033 | two component sensor kinase | 1.41 | 4E-62 |
| *chvI* | Atu0034 | two component response regulator | 2.27 | 3.6E-95 |

**Table S2. Bacterial strains and plasmids.** Bacterial strains and plasmids used in this study are listed below

| **Strain or Plasmid** | **Relevant Genotype, Features or Characteristics** | **Source or Reference** |
| --- | --- | --- |
| **Source Plasmid** |  |  |
| pNTPS139 | Km^r^; Suicide vector containing oriT and sacB | D. Alley |
| **Deletion Plasmid** |  |  |
| pNTPS138∆*chvI* | Km^r^ Suc^s^; deletion plasmid for *chvI* | Tomlinson et al[1] |
| pNTPS138∆*rem* | Km^r^ Suc^s^; deletion plasmid for zZ*rem* | Figueroa-Cuilan et al[2] |
| pNTPS139∆*exoA* | Km^r^ Suc^s^; deletion plasmid for *exoA* | This Study |
| pNTPS139∆T6SSpro | Km^r^ Suc^s^; deletion plasmid for T6SSpro | This Study |
| ***E. coli* strains** |  |  |
| DH5α | Cloning strain | Life Technologies |
| S17-1 | Smr;RP4-2 TC::MU Km-Tn7; for plasmid mobilization | Simon et al^6^ |
| ***A. tumefaciens* strains** |  |  |
| C58 | Parent strain | Watson et al[3] |
| C58∆*exoR* | ∆*exoR* | Tomlinson et al[1] |
| C58∆*chvG* | ∆*chvG* | Heckel et al[4] |
| C58∆*chvI* | ∆*chvI* | Heckel et al[4] |
| C58∆*tetRA*::a-*att*Tn7 (WT) | Replacement of the *tetRA* locus with an artificial *att*Tn7 site | Figueroa-Cuilan et al[2] |
| C58∆*tetRA*::a-*att*Tn7 ∆*rem* | ∆*rem* | Figueroa-Cuilan et al[2] |
| C58∆*tetRA*::a-*att*Tn7 ∆*exoA* | ∆*exoA* | This Study |
| C58∆*tetRA*::a-*att*Tn7 ∆T6SSpro | ∆T6SSpro | This Study |
| C58∆*tetRA*::a-*att*Tn7 ∆*pbp1b1* | *∆pbp1b1* | Williams et al[5] |
| C58∆*tetRA*::a-*att*Tn7 ∆*pbp1b2* | *∆pbp1b2* | Williams et al[5] |
| C58∆*tetRA*::a-*att*Tn7 ∆*pbp1c* | *∆pbp1c* | Williams et al[5] |
| C58∆*tetRA*::a-*att*Tn7 *∆mtgA* | *∆mtgA* | Williams et al[5] |
| C58∆*tetRA*::a-*att*Tn7 *∆pbp1b1,∆pbp1b2* | *∆pbp1b1,∆pbp1b2* | Williams et al[5] |
| C58∆*tetRA*::a-*att*Tn7 *∆pbp1b1, ∆pbp1b2, ∆pbp1c* | *∆pbp1b1,∆pbp1b2,∆pbp1c (∆*3pbp*)* | Williams et al[5] |
| C58∆*tetRA*::mini-Tn7-GM-Plac-pbp1a | Mini-Tn7T-GM-Plac-pbp1a inserted into a-*att*Tn7 site | Williams et al[5] |
| C58∆*tetRA*::mini-Tn7-GM-Plac -pbp1a, ∆*pbp1a* | Chromosome-based complementation of *∆pbp1a* with C58∆*tetRA*::mini-Tn7-GM-Plac-pbp1a allowing depletion of PBP1a under control of the lac promoter | Williams et al[5] |
| C58 ∆*tetRA*::mini-Tn7-GM-Plac -pbp1a, ∆*pbp1a,* ∆*rem* | ∆*rem* in PBP1a depletion background | This Study |
| C58 ∆*tetRA*::mini-Tn7-GM-Plac -pbp1a, ∆*pbp1a,* ∆*exoA* | ∆*exoA* in PBP1a depletion background | This Study |
| C58 ∆*tetRA*::mini-Tn7-GM-Plac -pbp1a, ∆*pbp1a,* ∆T6SSpro | ∆T6SSpro in PBP1a depletion background | This Study |
| C58 ∆*tetRA*::mini-Tn7-GM-Plac -pbp1a, ∆*pbp1a,* ∆*chvI* | ∆*chvI* in PBP1a depletion background | This Study |
| C58∆*tetRA*::a-*att*Tn7 *∆pbp1b1, ∆pbp1b2, ∆pbp1c* | *∆pbp1b1,∆pbp1b2,∆pbp1c (∆*3pbp*)* | Williams et al[5] |
| C58∆*tetRA*::mini-Tn7-GM-Plac -pbp1a, ∆*pbp1a* | Chromosome-based complementation of *∆pbp1a* with C58∆*tetRA*::mini-Tn7-GM-Plac-pbp1a allowing depletion of PBP1a under control of the lac promoter | Williams et al[5] |
| ***S. meliloti* strains** |  |  |
| Rm2011 | Wild type, Str^r^ | Casse et al[6] |
| Rm2011 *rgsP*-*egfp*, *∆*5pbp | Rm2011 *rgsP*-egfp carrying markerless deletions of *mrcA2, mcrB, pbp, pbpC* and SMc02856 *∆*5pbp | Williams et al[5] |
| Rm2011 *rgsP*-egfp *mrcA*1 depletion | Rm2011 *rgsP*-egfp carrying markerless deletion of *mrcA1*, curable complementation plasmid pGCH14-*mrcA1*, and pSRKKm as a source of *lacI* to cure pGCH14-*mrcA1*, Gm^r^ Km^r^ | Williams et al[5] |
| ***C. crescentus* strains** |  |  |
| NA1000 | Wild type | Hallez Lab |

**Table S3. Synthesized DNA primers.** The sequences of primers used to construct plasmids and strains in this study are listed below. All primers were ordered from IDT.

| **Synthesized DNA** | **Sequence (5’ – 3’)** |
| --- | --- |
| **Primers for deletion vectors in *A. tumefaciens*** |  |
| *exoA* P1 Forward SpeI | GCACACTAGTCGAGATCATCCTGCG |
| *exoA* P2 Reverse | AAGCTTGGTACCGAATTCAAGACCTTCCATGATTTG |
| *exoA* P3 Forward | GAATTCGGTACCAAGCTTAAAGGCTGTCTCATGACC |
| *exoA* P4 Reverse BamHI | CTGTCCTAGGCTTCCATCCTGAGAAGCG |
| *exoA* P5 Forward | TGCTGGTGACGAGTTCTCCGG |
| *exoA* P6 Reverse | AGTACCTGCACCACGCGG |
| *rem* P5 Forward | CATTATTTCCACGGCGAAAACTTCACCTC |
| *rem* P6 Reverse | GACCCGTGAAGCCATTGACGAC |
| T6SSpro P1 Forward SpeI | GCACACTAGTGCCTCTCCTGAACTTGTCAGC |
| T6SSpro P2 Reverse | AAGCTTGGTACCGAATTCATGTCGCATATCGATCTCAATCGCC |
| T6SSpro P3 Forward | GAATTCGGTACCAAGCTTTTGGATACACAGCATGTTAAAAG |
| T6SSpro P4 Reverse BamHI | CTAGCCTAGGGCTATCCGGTACAGTTCTTCG |
| T6SSpro P5 Forward | CGAGGTTCAGCAGGCAGACATTG |
| T6SSpro P6 Reverse | GCTTTCATCGGTGCCCGC |
| *chvI* P5 Forward | CGGCAGCAGGTAGTTCAGCAC |
| *chvI* P6 Reverse | CAGTGACAACACGATATTGACCAGCG |

**4) Supplementary Methods**

**Western blot analysis.** Cells were grown to exponential phase (OD = 0.5-0.8) in ATGN and then pelleted by centrifugation. Pellets were resuspended in 1X loading buffer and passed through a 20G needle to lyse the cells. 20 µL of each sample was loaded into each of three 4-20% Bis-Tris GenScript *SurePAGE* gels. BlueStain2 Protein ladder was loaded into the first well. Gel electrophoresis ran on ice at 100V for 90 minutes. After electrophoresis, one gel was Coomassie stained at room temperature with gentle shaking for 10-15 minutes. The gel was rinsed gently with water and then destained (H_2_O, methanol, acetic acid in a ratio of 50/40/10 v/v/v) at room temperature with gentle shaking for 10-15 minutes. The proteins in the remaining gels were transferred to an immobilon-FL transfer membranes cut to the size of the gel. Gel and membrane were secured and kept soaked with transfer buffer (25 mm Tris-Cl, 192 mm glycine, 20% (v/v) methanol) on ice overnight. Membranes were transferred to a solution of TBS + 0.05% Tween 20 with 4 µL of 1:1000 dilution of either anti-TssB or anti-Hcp and incubated at room temperature for 1 hour. Membranes was washed 3 times with fresh TBS + 0.05% Tween 20 for 5 minutes each. Immediately after wash steps, membranes were transferred to TBS + 0.05% Tween 20 with 4 µL of 1:1000 dilution of anti-rabbit HRP goat IgG as a secondary antibody and gently shaken for 1 hour. Membranes were washed 3 times with fresh TBS + 0.05% Tween 20 and then imaged. SuperSignal West Femto Maximum Sensitivity Substrate was used as a HRP substrate and imaging was done using a BioRad ChemiDoc Imager.
